# Synaptic architecture of leg and wing premotor control networks in *Drosophila*

**DOI:** 10.1101/2023.05.30.542725

**Authors:** Ellen Lesser, Anthony W. Azevedo, Jasper S. Phelps, Leila Elabbady, Andrew Cook, Durafshan Sakeena Syed, Brandon Mark, Sumiya Kuroda, Anne Sustar, Anthony Moussa, Chris J. Dallmann, Sweta Agrawal, Su-Yee J. Lee, Brandon Pratt, Kyobi Skutt-Kakaria, Stephan Gerhard, Ran Lu, Nico Kemnitz, Kisuk Lee, Akhilesh Halageri, Manuel Castro, Dodam Ih, Jay Gager, Marwan Tammam, Sven Dorkenwald, Forrest Collman, Casey Schneider-Mizell, Derrick Brittain, Chris S. Jordan, Thomas Macrina, Michael Dickinson, Wei-Chung Allen Lee, John C. Tuthill

## Abstract

Animal movement is controlled by motor neurons (MNs), which project out of the central nervous system to activate muscles. MN activity is coordinated by complex premotor networks that allow individual muscles to contribute to many different behaviors. Here, we use connectomics to analyze the wiring logic of premotor circuits controlling the *Drosophila* leg and wing. We find that both premotor networks cluster into modules that link MNs innervating muscles with related functions. Within most leg motor modules, the synaptic weights of each premotor neuron are proportional to the size of their target MNs, establishing a circuit basis for hierarchical MN recruitment. In contrast, wing premotor networks lack proportional synaptic connectivity, which may allow wing steering muscles to be recruited with different relative timing. By comparing the architecture of distinct limb motor control systems within the same animal, we identify common principles of premotor network organization and specializations that reflect the unique biomechanical constraints and evolutionary origins of leg and wing motor control.

## Introduction

All motor behaviors, from simple postural reflexes to complex locomotor navigation, are produced by patterns of electrical activity in MNs, which extend axons from the central nervous system to excite muscles throughout the body (Sherrington, 1906). A single MN and the muscle fibers it innervates comprise a motor unit. Animals achieve a remarkable diversity of behaviors by recruiting motor units in different combinations and sequences. Previous work has used analysis of limb biomechanics, movement kinematics, and electrophysiological recordings to identify these patterns of motor unit coordination (Lobato-Rios et al., 2022; Marshall et al., 2022; Ting and Macpherson, 2005; Tresch et al., 2002); however, the range of possible combinations of motor unit activations is ultimately constrained by the anatomy and connectivity of premotor neural circuits (Hodson-Tole and Wakeling, 2009). Thus, determining the wiring logic of premotor circuits can provide fundamental insight into how nervous systems coordinate motor units to accomplish diverse motor output.

Technical advances in reconstructing neurons and identifying synapses from volumetric electron microscopy (EM), or “connectomics”, have recently made it possible to analyze synapse-resolution circuit diagrams. There are now connectomes for the nematode *C. elegans* (Cook et al., 2019; Witvliet et al., 2021) and the larval fruit fly, *Drosophila melanogaster* (Ohyama et al., 2015; Winding et al., 2023) which provide insight into premotor control of peristaltic locomotion (Wen et al., 2012; Zarin et al., 2019). In addition to multiple connectomes of the adult fly brain (Scheffer et al., 2020; Zheng et al., 2018), there also exist two EM volumes of the adult *Drosophila* ventral nerve cord (VNC), from a female (<u>F</u>emale <u>A</u>dult <u>N</u>erve <u>C</u>ord, FANC; Phelps et al., 2021) and male fly (Takemura et al., 2023). The fly VNC functions like the vertebrate spinal cord to sense and move the fly’s legs and wings (Court et al., 2020), making these connectome datasets the first in which it is possible to analyze limb premotor circuits. We recently combined reconstruction of leg and wing MN dendrites in the FANC dataset with genetic tools and x-ray holographic nanotomography to map the muscle targets of MN axons to the level of individual muscle fibers (Azevedo et al., 2022). This high-resolution atlas of MN to muscle connectivity now makes it feasible to compare how premotor circuits are organized to control two evolutionarily distinct limbs, the leg and wing.

Flies use their legs to perform many behaviors, including locomotion, grooming, aggression, and courtship (Tuthill and Wilson, 2016). The position of the distal tip of the fly’s leg is determined by seven mechanical degrees of freedom, specified by the biomechanics of the five joints within each leg. The most proximal joint (thorax-coxa) has three degrees of freedom, while the other joints have one (Lobato-Rios et al., 2022). Most leg joints are actuated by more than two muscles, e.g. two muscles in the femur contribute to tibia flexion and one antagonist muscle controls tibia extension. In total, the fly’s front leg contains 18 muscles (Miller, 1950), collectively innervated by 69 MNs (Azevedo et al., 2022). Most leg muscles are innervated by multiple MNs, i.e. they are composed of 2-8 motor units, with MNs innervating distinct fibers within the target muscle (Azevedo et al., 2022). Unlike some other insect species, flies lack GABAergic MNs (Schmid et al., 1999; Witten and Truman, 1998). In many animals, a motor pool refers to groups of motor units that are recruited together, and can include motor units across multiple muscles. The dynamic firing patterns of motor pools, including antagonist pools, control joint torque by causing muscle contractions that interact with complex, passive musculoskeletal biomechanics.

In many animals, from flies to humans, MNs within a motor pool typically fire in a stereotyped order, or recruitment hierarchy, which helps to smoothly increase muscle force in a graded manner (Gabriel et al., 2003; Henneman et al., 1965a; Hill and Cattaert, 2008; Marshall et al., 2022; McLean and Dougherty, 2015; Milner-Brown et al., 1973; Sasaki and Burrows, 1998). Since the pioneering work of Henneman (Henneman, 1957), it has been theorized that the MN recruitment hierarchy is established by correlations in physiological and neuromuscular properties that are collectively referred to as the size principle. The size principle posits that if a common synaptic input excites a motor pool, MNs with the highest input resistance. i.e., small MNs that innervate slow-twitch fibers, will depolarize and fire first. As the strength of the common input increases, larger MNs that innervate fast-twitch fibers will then be recruited, establishing a recruitment order based on MN size (Henneman et al., 1974). The precise synaptic architecture of limb premotor circuits that is responsible for implementing the size principle has remained a mystery since the phenomenon was first described in cat leg muscles over 60 years ago (Henneman, 1957). Recent work in adult zebrafish has revealed that separate populations of interneurons synapse onto small/slow vs. large/fast MNs, raising the question of how common input to a motor pool contributes to the recruitment hierarchy (Pallucchi et al., 2024; Song et al., 2020). In addition, modern muscle recordings in primates demonstrate that the correlations in motor unit firing rates can flexibly change with the demands of the motor task, in contrast to stereotyped recruitment that is the foundation of the size principle (Marshall et al., 2022).

The fly wing is biomechanically and evolutionarily distinct from the leg (Grimaldi and Engel, 2005; Hörnschemeyer, 2002). One key difference is that wing MNs do not possess conventional MN recruitment hierarchies. Three separate groups of muscles with specialized physiological properties and functions control each wing (Melis et al., 2024; Miyan and Ewing, 1985). One group of large muscles, called *indirect* because they do not directly attach to the wing hinge, span the thorax orthogonally, forming a pair of antagonist muscles that provide the power to flap the wings. The indirect muscles are innervated by 12 MNs per side. They are asynchronous, stretch-activated muscles, so their contractions arise at the level of muscle biomechanics, rather than from neural input (Coggshall, 1978; Pringle, 1949). Another group of muscles, known as *tension* muscles, modify the stiffness of the thorax to modulate the power produced by the indirect muscles (Nachtigall and Wilson, 1967; Pringle, 1957). Forces from the resonant oscillations of the thorax are transmitted through the intricate linkages of the wing hinge to produce the trajectory of the wing stroke (Lehmann and Dickinson, 2001, 1997). The third group of muscles are directly attached to cuticle thickenings, or sclerites, that make up the wing hinge (Boettiger and Furshpan, 1952; Miyan and Ewing, 1985; Trimarchi and Schneiderman, 1994; Williams and Williams, 1943). The actions of these *direct steering* muscles create subtle deviations in the wing trajectory that modify aerodynamic forces for flight control and acrobatic maneuvers (Dickinson and Tu, 1997). Each direct muscle is innervated by a single MN, so wing steering motor units are not organized into conventional motor pools. Although past work has revealed mechanisms of flight motor control at the level of muscles and single MNs (Balint and Dickinson, 2001; Dickinson et al., 1993; Heide, 1983, 1975; Lehmann and Bartussek, 2017; Lindsay et al., 2017; Melis et al., 2024; O’Sullivan et al., 2018; Whitehead et al., 2022) comparatively little is known about wing premotor circuits within the VNC that coordinate MNs to control flight and other wing-related behaviors. Here, we use connectomics to comprehensively reconstruct premotor synaptic input to leg and wing MNs, allowing us to compare similarities and differences in neural control of joints with distinct biomechanics.

## Results

We reconstructed the anatomy and synaptic connectivity of all premotor neurons (preMNs) that synapse onto MNs that control either the left wing or the left front leg in the FANC dataset (**Figure 1a-d**; Azevedo et al., 2022). We focus here on the fly’s front leg, which is specialized for tactile exploration and possesses a greater range of motion than the middle and rear legs (Grabowska et al., 2012). The resulting premotor connectome consists of 69 leg MNs that receive 212,190 synapses from 1,546 preMNs, and 29 wing and thorax MNs that receive 144,668 synapses from 1,784 preMNs. MNs have tortuous dendritic arbors and possess among the largest cell bodies in the *Drosophila* nervous system (**Figure 1e-f**). On average, each MN receives 3,641 input synapses from 188 preMNs (using a 3-synapse threshold; quantification for each MN in **Extended Data Figure 1a-h**) and each preMN synapses onto 6 MNs (7.2±7.4 sd. for leg preMNs, 5.1±3.1 for wing preMNs).

**Figure 1.**
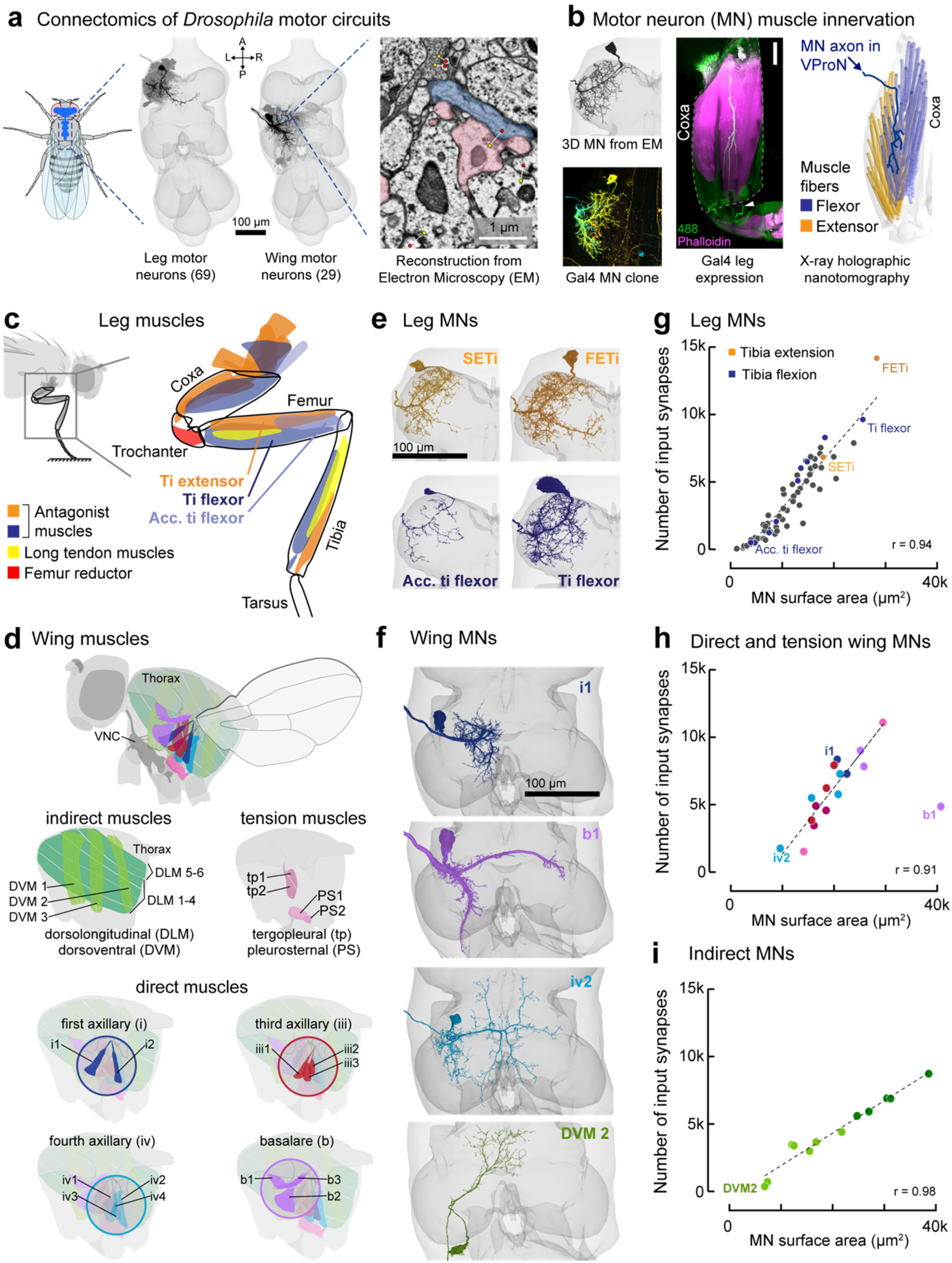
Reconstruction of synaptic inputs to identified leg and wing MNs in *Drosophila*. **a**, Automated neuron segmentation and synapse prediction in a serial-section electron microscopy (EM) volume of an adult female VNC. The VNC contains the cell body and dendrites of MNs controlling the leg (left, N = 69) and the wing (right, N = 29). Axons extend out of the VNC to innervate muscles. Example EM image shows automated synapse predictions, with presynaptic sites labeled by yellow dots and postsynaptic sites by red. **b**, Identification of muscle targets of each leg MN (Azevedo et al., 2022). Briefly, reconstructed EM neurons (top left) were compared with X-ray tomography of the leg (right) and light-microscope images of genetically identified MNs in the VNC (bottom left, Meissner et al, 2023) and in the leg (middle; GFP and cuticle autofluorescence in green, phalloidin in magenta, scale bar is 50 µm). Here, a clone of a trochanter flexor MN labeled by VT063626-Gal4 (Meissner et al, 2023), with axon expression in the trochanter flexor muscle in the coxa. The axon travels in the ventral prothoracic nerve. White arrowhead indicates a sensory axon in the prothoracic leg nerve, illustrating that not all Gal4 lines are specific for MNs. **c**, Schematic of the 18 muscles controlling the front leg. **d**, Flight musculature, separated according to muscle physiology and/or insertion anatomy. **e**, Reconstructed MNs that extend and flex the tibia. **f**, Reconstructed wing MNs. **g**, Leg MN input synapses scale linearly with surface area, such that synapse density is 0.45 synapses μm^-2^ (r=0.94, p<10^-33^). **h**, Input synapses vs. surface area for direct and tension wing MNs (density=0.41 synapses μm^-2^, r=0.90, p<0.10^-3^, excluding the outlier b1 MN, see Methods) **i,** Input synapses vs. surface area for indirect wing MNs (density=0.24 ±0.02 synapses μm^-2^, r=0.98, p<10^-7^).

In all limbed animals studied to date, measurements of MN size, such as soma volume, axon diameter, and dendritic surface area, have been found to correlate with the number and physiology of the muscle fibers the MN innervates and with the amount of force the motor unit produces (Burrows, 1996; Cullheim, 1978; Hill and Cattaert, 2008; Hoyle, 1983; Kernell, 2006; Mcphedran et al., 1965; Monster and Chan, 1977; Wuerker et al., 1965). When we reconstructed dendrites and somas of fly leg and wing MNs, we found that MN surface area and volume ranged over 40-fold (**Figure 1g, h; Extended Data Figure 1i-k**). This variation in MN size is consistent with the range in MN morphology in the cat (Cullheim et al., 1987; Kernell and Zwaagstra, 1989). Note, the somas of insect neurons do not typically receive synaptic input and thus may not serve the same important role in neuronal processing as they do in vertebrate neurons.

We found that the total number of MN input synapses is highly correlated with MN surface area, but that the relative densities are different for MNs that innervate different types of muscles. Leg MNs have a synapse density of ∼0.45 synapses per μm^2^ (**Figure 1g**; r=0.94, p<10^-33^). This density is approximately 3X more synapses per unit area than reported for cat MNs (Örnung et al., 1998). Consistent with the size principle, we found that each leg muscle is innervated by MNs of different sizes (**Extended Data Figure 1i**), so larger MNs receive proportionally more synapses. MNs that innervate direct and tension wing MNs have a similar density to leg MNs, 0.41 synapses μm^-2^ (**Figure 1h**; r=0.90, p<0.10^-3^, excluding the outlier b1 MN, see **Methods**). Indirect MNs have a lower synapse density, with ∼0.24 synapses μm^-2^ (**Figure 1i**; r=0.98, p<10^-7^). In summary, our quantification of synaptic input to MNs identified previously unknown correlations between MN size and number of input synapses across both leg and wing MNs.

If MN recruitment were dictated entirely by the magnitude of premotor input (**Figure 1h**), then these patterns of proportional connectivity would cause a common input to first recruit the largest, fastest MNs. However, MNs controlling the fly tibia follow a conventional recruitment hierarchy during spontaneous movements, with the smaller MNs in a pool firing prior to larger MNs (Azevedo et al., 2020). Indeed, the largest MNs in the tibia motor pool rarely spike during behavior (Azevedo et al., 2020). Thus, the intrinsic electrical properties of the largest MNs must compensate for the higher number of input synapses to establish a recruitment hierarchy that follows the size principle.

### Local neurons dominate synaptic input to both leg and wing MNs

We next set out to dissect the sources of input to MNs and to compare the quantitative structure of synaptic connectivity for leg and wing premotor networks. We classified preMNs into five morphological classes: descending (from the brain), sensory, ascending (to the brain), intersegmental (from other VNC neuropils), and local neurons (**Figure 2a-c**). **Figure 2d-e** show the entire leg and wing premotor connectivity matrices: preMNs are rows, MNs are columns, and the color of each element indicates the number of synapses between the two cells. We sorted leg MNs from proximal to distal, according to the location of the muscle fibers they innervate (Azevedo et al., 2022). For the wing, we ordered indirect wing MNs first, followed by tension MNs, and then direct MNs, sorted by the wing sclerite upon which they insert (**Figure 2e**). We sorted preMNs of each class by their top MN targets (per muscle), and then by total synapses onto MNs (the sum of each row).

**Figure 2.**
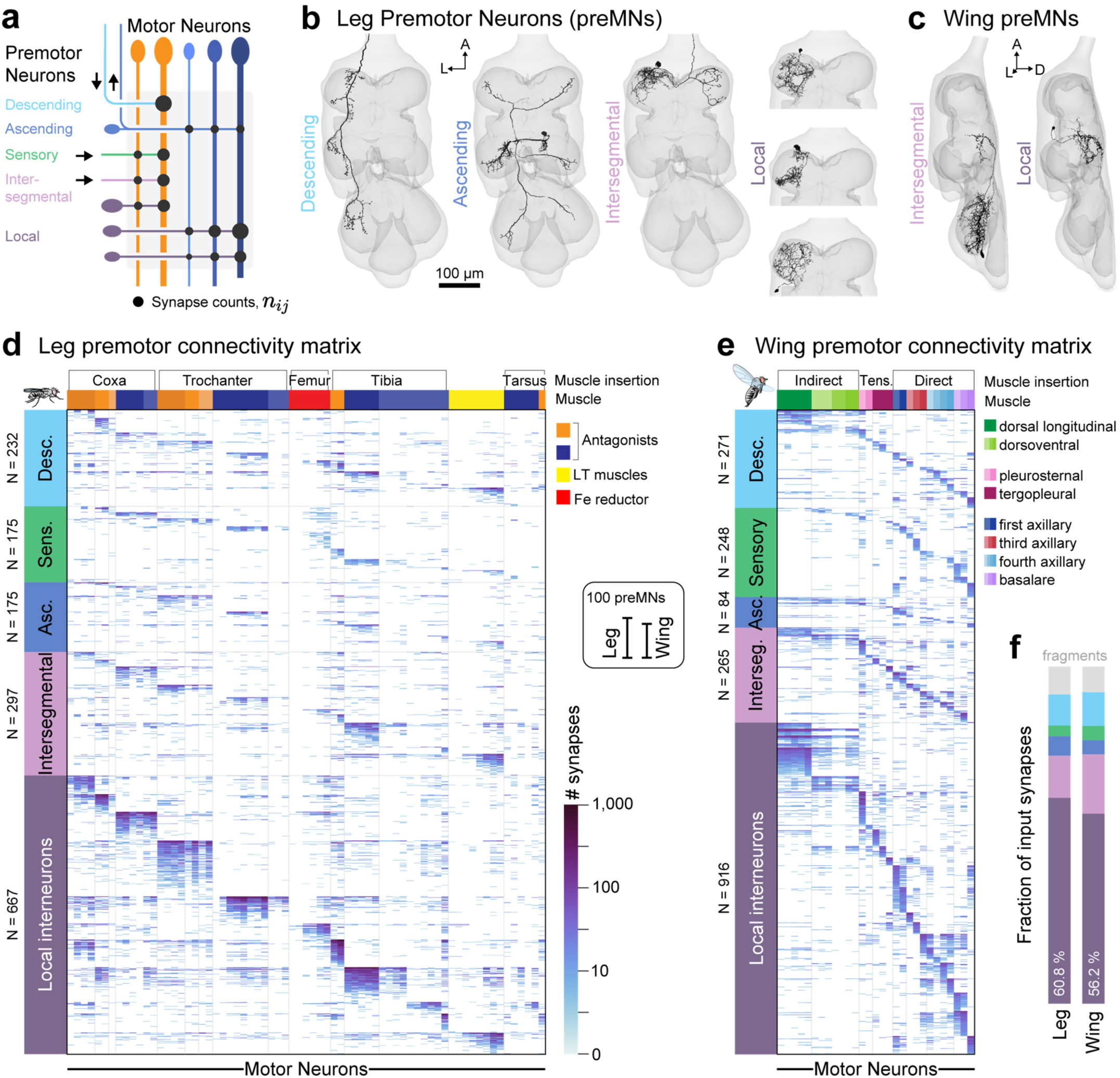
The majority of synaptic input to MNs is from local interneurons. **a,** Schematized premotor connectivity matrix. PreMNs (rows) are colored according to cell class. MNs (columns) are grouped by their distinct muscle targets (orange and blue). **b,** Examples of descending, ascending, intersegmental, and local leg preMNs. Intersegmental leg preMNs receive input from other neuromeres. **c,** Example intersegmental and local wing preMNs. Intersegmental wing preMNs receive input from leg neuromeres. **d,** Premotor connectivity matrix for the left T1 leg. The color of each element indicates the log10(n+1) of the number of synapses from each preMN (row) onto each MN (column). Note that the log scale makes differences appear smaller than they are. Gray vertical lines delineate muscle targets, from proximal to distal, indicated by color (top). PreMNs are ordered: 1) by cell class; 2) by MN targets; 3) by total synapses onto all MNs. Connectivity matrix only shows synapses from fully proofread neurons; 19,349 synapses from fragments account for 8.3% of the left front leg premotor connectome (see Methods). **e,** Same as **d** for wing MNs. 11,966 synapses from fragments account for 7.6% of the left wing premotor connectome. **f,** Fraction of total synaptic input from each cell class onto leg or wing MNs. Fragments do not belong to any class, because they have not been attached to a primary neurite via proofreading.

We found that the predominant source of synaptic input to both leg and wing MNs is local VNC neurons (**Figure 2d-f**, **Extended Data Figure 1c-d**). Although previous work in the fly has focused on the contributions of descending neurons (DNs) to fly motor control (Bidaye et al., 2014; Cheong et al., 2024; Feng et al., 2020; Namiki et al., 2018; Schnell et al., 2017), we find that less than 10% of synapses to each leg and wing MN comes from DNs (mean = 9.1 ±4.2% sd. for leg MNs and 11.2 ±6.5% for wing MNs). Local preMNs, however, comprise 43% of all leg preMNs but 63.4±9% of synaptic input to each MN (**Extended Data Figure 1j**). These proportions suggest that most descending signals are routed through local premotor circuits, which then coordinate groups of MNs.

We also observed input connectivity patterns that reflect specializations of specific MNs. For example, some wing steering MNs fire tonically during flight (Lindsay et al., 2017; Melis et al., 2024), at rates as high as wingbeat frequency (Heide and Götz, 1996) We found that the MNs to four tonic muscles were the only cells with >10% of synaptic input from sensory neurons (iii3=18.5%, b1=17.3%, b3=13.5%, i2=11%; **Extended Data Figure 1d**). This particular organization may reflect the importance of wingbeat-synchronous proprioceptive feedback in tuning the activation phase of these muscles (Fayyazuddin and Dickinson, 1998), a feature that is known to modulate their wing stroke trajectory on a stroke-by-stroke basis (Tu and Dickinson, 1994).

In summary, we found quantitative similarities in the proportions of synapses to leg and wing MNs from different classes of preMNs. We next sought to understand how the structure of this premotor connectivity relates to the organization of leg and wing muscles.

### Hierarchical clustering of MN input connectivity reveals patterns of common synaptic input

Common synaptic input to MNs within a motor pool has been proposed to underlie hierarchical MN recruitment by supporting graded production of muscle force to control a single joint (De Luca and Erim, 1994; Henneman et al., 1974). On the other hand, common input to MNs that actuate different joints is thought to underlie coordinated firing patterns during complex multi-jointed movements, termed a muscle synergy (Ting and McKay, 2007; Tresch et al., 2002). Because we previously mapped the muscle innervation of each leg and wing MN from the FANC connectome (Azevedo et al., 2022), we were able to analyze the relationship between common premotor input to MNs and the identity of the muscles they control.

To quantify common input to MNs, we first compared pairs of MNs based on the number of synapses from preMNs (**Figure 2d, e**). We used cosine similarity, an established metric for comparing the similarity of vectors (Li et al., 2020; Matsliah et al., 2023). Cosine similarity is the dot product of the normalized inputs (L2-norm of the column vectors; **Figure 3a-d**). Normalization turns the columns into unit vectors in N-dimensional preMN space (N=1,546 leg preMNs or N=1,784 wing preMNs), which facilitates comparing MNs that vary in the total number of synapses they receive (**Figure 3b**). Values of high cosine similarity indicate that two MNs receive the same synaptic weights from the same preMNs, i.e. the same number of synapses in proportion to total synaptic input (**Figure 3b**), suggesting a high likelihood of co-activation or co-inhibition. Low similarity scores indicate either that two MNs share few synaptic partners or that the relative input weights from common preMNs are different enough to pull the column vectors in different directions (**Figure 3c**). The matrices of pairwise cosine similarity for leg MNs (**Figure 3e**) and wing MNs (**Figure 3f**) revealed distinct clusters of MNs with high cosine similarity, reaching extremely high values (>0.9) for some pairs.

**Figure 3.**
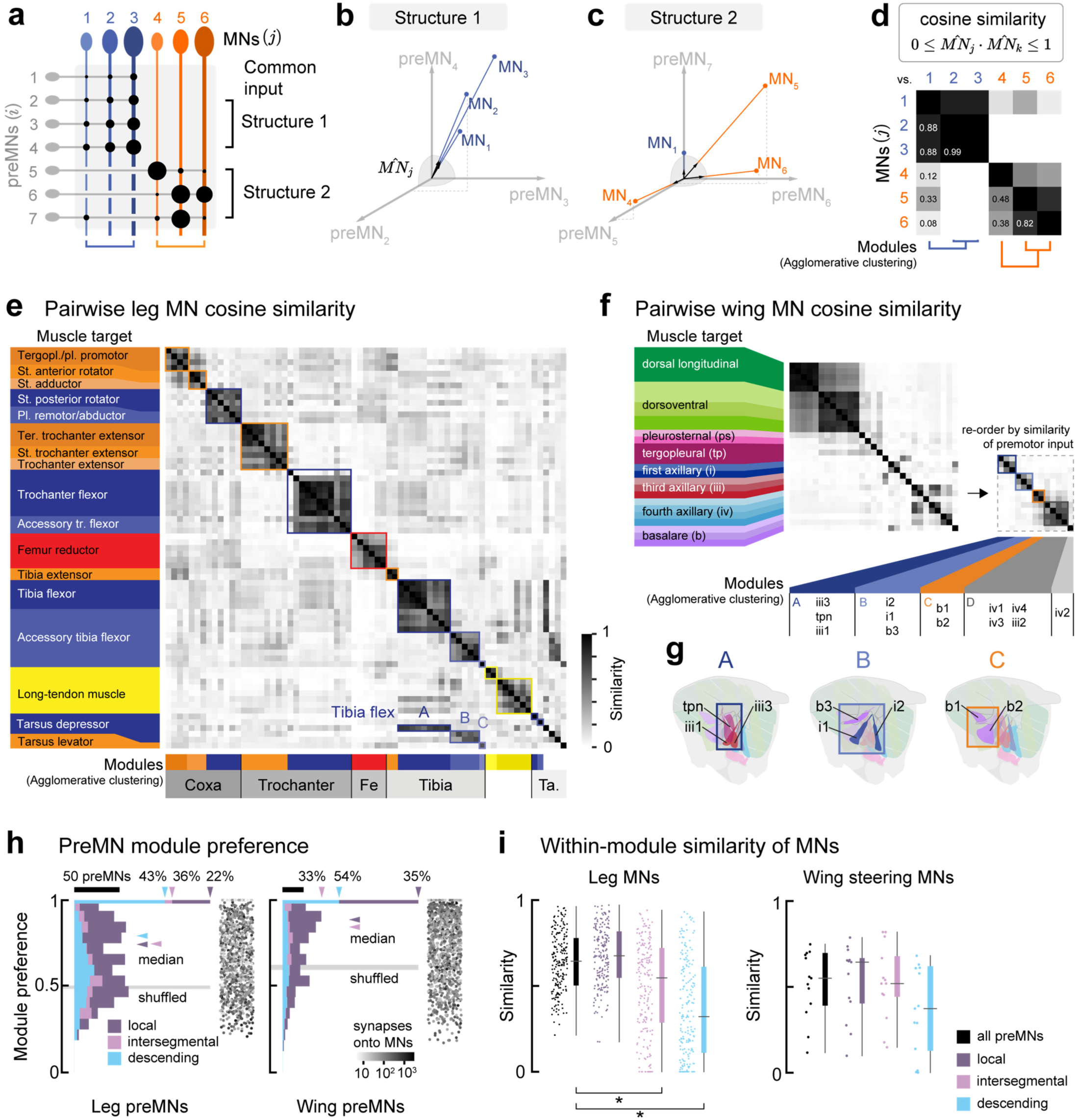
PreMNs preferentially target specific groups of MNs, forming motor modules. **a**, Schematic illustrating how cosine similarity depends on the structure of preMN input. Each MN can be thought of as a vector in a preMN space, with seven dimensions in this example. **b-c,** Cosine similarity is the dot product of the normalized column vectors in premotor space (black arrows) on the unit sphere (gray). **b**, If MNs (blue) receive the same normalized synaptic weights from the same preMNs, the pairwise similarity is near 1 (Structure 1). **c,** If MNs (orange) receive input from the same preMNs, but not the same weights, the cosine similarity will be non-zero, but less than 1 (Structure 2). **d,** Matrix of pairwise cosine similarity for MNs. Agglomerative clustering defines groups of MNs with high cosine similarity, which we call motor modules. **e,** Pairwise MN cosine similarity matrix for leg MNs. MNs are ordered as in Figure 2, from proximal to distal muscle targets. Muscle innervation of each MN is indicated at left. Motor modules are indicated below (see **Extended Data Figure 2a-f** for details). **f,** Pairwise similarity of wing MNs. Wing MNs are ordered by sclerite attachment as in Figure 2. Inset shows reordered direct steering MNs by module structure (defined in **Extended Data Figure 3a-d**). **g,** Steering modules are composed of muscles that attach to distinct sclerites. **h,** PreMN module preference (see Methods) for local (purple), intersegmental (pink), and descending (blue) leg preMNs, the three cell classes that account for most MN input. Vertical arrowheads indicate the fraction of preMNs that synapse onto a single module, i.e. module preference = 1. Horizontal arrowheads indicate the median module preference. Gray box is the range of median module preference when rows of the connectivity matrix are shuffled (N=10^4^ shuffles). Gray dots show total number of MN synapses (log-scale) for individual preMNs. Module preference is not strongly correlated with total MN synapses (All leg preMNs: Pearson’s r=-0.29, p<10^-30^; wing preMNs: r=-0.18, p<10^-14^). **i,** Pairwise similarity of MNs within modules, when considering all preMNs (black), or just a single class. Left: within-module leg MN similarity is lower when considering only intersegmental or descending preMNs (p<10^-24^, Kruskal-Wallis non-parametric one-way ANOVA. Conover’s post-hoc pairwise test with Holm correction for multiple comparisons: * - p<10^-6^). Right: within-module similarity of wing MNs (excluding indirect and tension muscle MNs) is generally lower than for leg MNs.

To identify groups of MNs that share common premotor input, we performed unsupervised agglomerative clustering on the cosine similarity matrices (**Figure 3d-f**, see **Extended Data Figures 2, 3** for details of clustering procedures). This ordering allowed us to group MNs according to the similarity of their premotor networks, irrespective of the muscles they innervate. We refer to each cluster as a “motor module”, because MNs within each cluster are likely to be recruited together due to their common preMN input.

#### Leg MNs

Most leg motor modules contained MNs that innervate the same or mechanically linked muscles that attach to the same joint (**Figure 3e**). This organization is illustrated by the two modules that control the trochanter. Seven MNs that innervate the trochanter flexor muscle all cluster together, along with the MNs that innervate the accessory trochanter flexor, a separate muscle that actuates the same joint. The antagonist trochanter extensor module is composed of eight MNs that innervate three parts of the muscle with different origins and insertions on the same extensor tendon. The antagonist trochanter flexors and extensors have cosine similarity near zero, indicating they share few sources of input. The coxa muscles featured a different module organization, where the seven MNs innervating the four muscles that insert on its anterior aspect separate into two distinct modules. The coxa has three degrees-of-freedom, so the two promotor muscles (4 MNs) and the rotator and adductor muscles (3 MNs) are not strict synergists and thus may not share a common recruitment hierarchy, similar to the human shoulder (Wickham and Brown, 1998). These examples of the trochanter and coxa illustrate our general finding that the composition of most leg motor modules is consistent with the size principle model that MNs within a motor pool receive common input from preMNs.

Clustering MNs by their preMN input also revealed some unexpected modules that indicate novel and specialized functions. First, the accessory tibia flexor MNs separate into three different clusters (**Figure 3e**). This finding is not necessarily inconsistent with the motor pool concept, as individual muscle fibers of the accessory tibia flexor muscle insert on the tibia through individual thin tendons, rather than inserting onto a single large tendon like the tibia flexor muscle (Azevedo et al., 2022). Subdividing tibia modules may improve dexterity by allowing each muscle fiber some mechanical independence, as has been observed in the locust tibia (Hoyle, 1955; Sasaki and Burrows, 1998). Second, MNs that innervate tarsus control muscles receive little common input. Instead, four of the six tarsus depressor MNs share common premotor input with each of the three clusters of tibia flexor modules, respectively. Thus, the most distal segments may be controlled by shared premotor input that produces co-contraction of tarsus and tibia muscles.

#### Wing MNs

Unsupervised agglomerative clustering of the wing premotor network revealed two subdivisions of wing MN modules that map onto the two anatomically and physiologically distinct muscle systems: indirect MNs and direct/tension MNs. Further, the MNs of the indirect muscles clustered into two groups, corresponding to the dorsolongitudinal muscles (DLMs, 5 MNs) and dorsoventral muscles (DVMs, 7 MNs). This organization is consistent with the antagonistic function of the DLMs and DVMs, which power the downstroke and upstroke, respectively. Notably, the two indirect modules also receive substantial shared input (similarity = 0.5 ±0.07). This similarity makes sense because the indirect muscles contract asynchronously from MN spikes; indirect MNs all spike at the same frequency during flight, though their timing is offset from each other (Coggshall, 1978; Hürkey et al., 2023; Koenig and Ikeda, 1983). Thus, while power MNs receive a high degree of common preMN input, these input patterns are unlikely to contribute to the establishment of a recruitment hierarchy.

Pairs of wing tension MNs, on the other hand, exhibit low similarity (**Figure 3f**, mean = 0.13 ±0.10), suggesting they do not receive substantial common input. These five MNs have large numbers of input synapses, on par with the largest leg MNs (**Extended Data Figure 1b**). However, tension MN synapses are largely from non-overlapping populations of preMNs. These distinct preMN populations may represent parallel premotor pathways for independent control. Interestingly, one tension MN, named “tpn” as it innervates both the tp1 and tp2 fibers of the tergopleural muscle (O’Sullivan et al., 2018), displayed higher similarity with two direct MNs than with other tension MNs.

Unlike most leg modules, we observed high similarity between MNs innervating direct (i.e., steering) muscles that attach on different sclerites within the wing hinge (**Figure 3f, g**). Each direct muscle is innervated by a single MN and inserts on its sclerite at a different orientation to exert force in different directions. Each wing sclerite is equipped with at least one tonic muscle and one phasic muscle, which permits both constant trimming of the hinge mechanics as well as additional activation during rapid maneuvers (Lindsay, et al., 2017). Based on this organization, MNs controlling muscles attached to the same sclerite may need to be recruited independently, rather than all together. Consistent with this prediction, clustering delineated four distinct modules of direct MNs (plus the one tension MN) that receive common premotor input (**Figure 3f**, inset). The compositions of some of these modules are also supported by prior physiological recordings, such as anti-correlations between i1 and iii1 and correlations between b1 and b2 spiking within a wing stroke (Heide, 1975). More recent work found correlations in muscle calcium signals that perfectly match this premotor module structure (Melis et al., 2024). Thus, both neuromuscular recordings and clustering by preMN synaptic connectivity suggest that wing steering MNs form synergies that are coordinated by dedicated premotor pathways.

### PreMNs preferentially synapse onto specific motor modules

Unsupervised clustering revealed modularity of MNs due to shared preMN input (**Figure 3e, f**). We next sought to understand how preMN synaptic connectivity patterns give rise to this modular structure. Do all preMNs synapse predominantly onto a single module, or do some preMNs connect broadly across modules? For each preMN, we computed the “module weights”, defined as the number of synapses a preMN makes in each module relative to its total number of synapses onto MNs. We defined “module preference” as the highest module weight of each preMN. The median module preference is 0.80 (interquartile range (iqr) =0.45) for leg preMNs and 0.90 (iqr=0.33) for wing preMNs. For leg modules that have clear antagonist modules, we found that the module weight for the antagonist module is low (mean of 0.012±0.001 (s.e.m.) vs. 0.02±6<10^-4^ for module weights for all non-preferred modules, p<10^-6^ Mann-Whitney U test). When we shuffled the preMN synapse counts, the median module preference decreased to 0.50 (**Figure 3h**), whereas the mean antagonist module weight increased ∼10-fold to 0.11±0.06 (s.e.m., N=10,000 shuffles). The preference for a specific motor module thus varies across preMNs, but most leg and wing preMNs exhibit a preference for a specific motor module that is higher than would be expected from random wiring. Illustrating the strength of this preference, preMN synapses onto a preferred module account for 62.2% of synapses onto leg MNs and 75.7% of synapses onto wing MNs (**Extended Data Figure 4b, f**).

Even though preMNs tend to synapse on MNs in a single preferred module, this does not always result in high MN cosine similarity. For example, the cosine similarity of wing steering MNs is never as high as leg MNs (**Figure 3i**), even though wing preMNs have high module preference (**Figure 3h**). One way this could arise is depicted in **Figure 3c**: both network structures contain preMNs that preferentially target a single module, but Structure 2 results in lower cosine similarity because the synaptic weights of its preMN inputs are more variable across their MN targets. The cosine similarity of leg MNs is also lower if it is calculated using only the connections from intersegmental or descending preMNs (**Figure 3i, Extended Data Figure 4c**). By contrast, for both leg and wing MNs, local preMN connectivity results in a similar distribution of cosine similarity for MNs in the same module (**Extended Data Figure 4d, h, and j**). Together, this analysis suggests that the connectivity of local preMNs, as compared to other cell classes, is the primary driver of MN modularity. It also suggests that connectivity of local preMNs is structured differently for leg and wing motor systems, a hypothesis we explore next.

### The structure of local preMN input to leg and wing motor modules reflects their unique biomechanics

We next analyzed the structure of local preMN input within motor modules. We first focused on the tibia, where it is well established that MNs are hierarchically recruited (from slow/small to large/fast) in flies and other insects (Azevedo et al., 2020; Burrows, 1996; Newland and Kondoh, 1997; Sasaki and Burrows, 1998). We quantified the structure of local preMN synaptic input onto MNs in their preferred module (**Figure 4a**). We defined the output weight, *w_ij_*, as the synapse count from preMN, *i*, onto MN, *j*, divided by the sum of the synapses onto the entire motor module. We found that each preMN provides the same output weight onto all the MNs within the extensor (**Figure 4b**) as well as flexor modules (**Figure 4c**). Specifically, the preMN output weights onto each MN are proportional to the overall synaptic input to each MN. This is true regardless of the total number of preMN synapses, which can range over 100-fold (**Figure 4b-c, Extended Data Figure 5a-c**). Shuffling the connections onto the most similar MNs lowered the mean cosine similarity, showing that the pattern of proportional synaptic weights is a key factor that determines MN input similarity (**Extended Data Figure 5d-g**). We observed this pattern in all leg motor modules with more than one MN.

**Figure 4.**
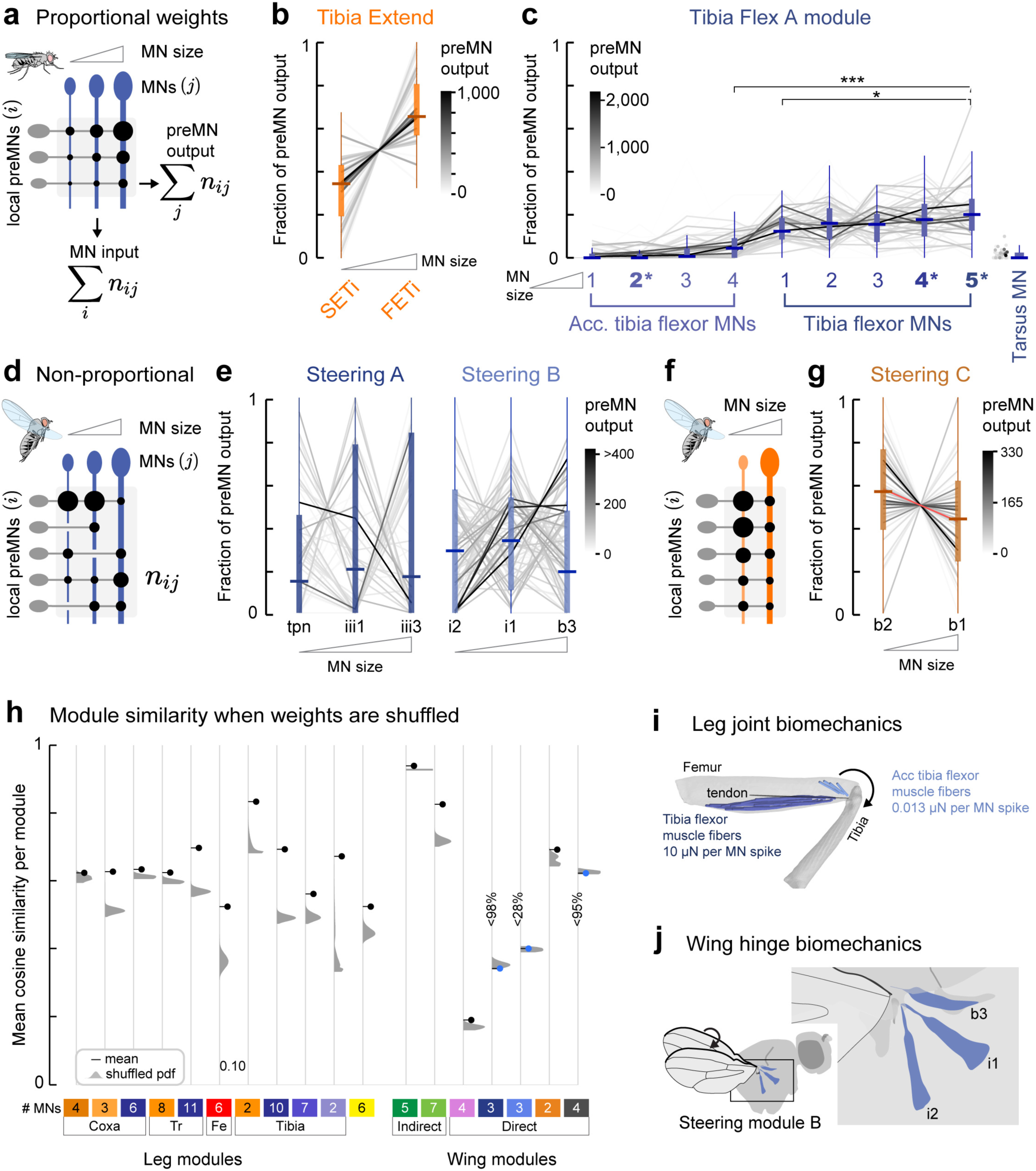
Local preMN connectivity differs between leg and wing steering motor modules. **a,** Illustration of proportional connectivity of module-preferring preMNs in leg modules. PreMN output is defined as the sum of a preMN’s synapses onto MNs in the module. **b,** Proportional connection weights from local preMNs that prefer the Tibia Extensor module onto the Slow (SETi) and Fast (FETi) tibia extensor MNs. Line hue indicates total number of synapses onto the module. **c,** Proportional connection weights from local preMNs that prefer the Tibia Flex A module onto the Tibia Flex A module. The module is composed of four accessory tibia flexor MNs, the five tibia flexor MNs, and the synergist tarsus MN. Tibia MNs are ordered by surface area. Asterisks (*) indicate MNs previously characterized using electrophysiology (Azevedo 2020). The fraction of preMN output onto the largest MN is significantly larger than onto smaller MNs (p<10^-4,^, Kruskal-Wallis non-parametric one-way ANOVA. Conover’s post-hoc pairwise test with Holm correction for multiple comparisons: *** - p<10^-28^, * - p=0.012). **d,** Wing steering motor modules lack proportional connectivity. **e,** Local preMN connections onto wing Steering Modules A and B. **f,** Local preMNs make more synapses onto the smaller b2 MN than the larger b1 MN. **g,** Same as **e** for Steering Module C. **h,** Average cosine similarity (black dots) compared to the average cosine similarity when synapse counts from local preMNs onto their preferred module are shuffled (gray probabilities density function (PDF), N=10,000 shuffles). The measured mean is larger than 99% of the PDF in all cases except for coxa promotors (left, larger than 98.3%), steering module B (62%), and steering modules A and D (larger than only 2% and 5% of shuffles, respectively). **i,** MNs in the Tibia Flex A module work together to produce torque on the tibia. Measurements of tibia force produced by single spikes in the 2* and 5* MNs from Azevedo et al., 2020. **j,** Schematic of muscles innervated by wing steering module B. Each muscle has a different origin and attachment to the wing hinge, suggesting co-contraction of muscles can have variable biomechanical effects.

When we plotted the preMN synaptic weights onto wing steering modules, we found a different, non-proportional structure (**Figure 4d-g**). Specifically, preMNs that contact each MN in a wing steering module do not tend to distribute their synapses in proportion to overall synaptic input. Even steering Module C, which contains only the b1 and b2 MNs, is characterized by wider variance of weights (**Figure 4g**) when compared with the pair of tibia extensor MNs (**Figure 4b**). In this case, the smaller b2 MN receives a greater proportion of input from the majority of common local preMNs.

To explore how the structure of preMN synaptic weights compares to a random connectivity scheme, we computed the average cosine similarity of MNs when synapse counts are shuffled within the module (**Figure 4h**). For most modules, shuffling lowered the mean cosine similarity, consistent with most modules having a proportional connectivity structure. By contrast, if the synapse counts for wing steering modules A or D were shuffled, the resulting average cosine similarity was almost always higher, suggesting that these non-proportional connectivity patterns are not random (**Figure 4h**). The proportional and non-proportional structures of leg and wing modules are also supported by principal component analysis (PCA) of the leg and wing modules (**Extended Data Figure 6a**).

In summary, the strength of connections from local preMNs to leg motor modules is proportional to total MN input, while local preMN input to wing steering motor modules is not. This difference explains why modules controlling the wing steering muscles have lower within-module similarity than leg modules (**Figure 3i, Extended Data Figure 4i**). The difference in preMN connectivity structure in wing modules may reflect specialized motor control strategies for the leg and wing, which possess distinct biomechanics (**Figure 4i-j**). Tibia flexor MNs in the same module act together to control flexion torque, but produce force per spike that ranges over three orders of magnitude (Azevedo et al., 2020). By contrast, the muscles that form each wing steering module have different origin and insertion points (**Figure 4j**). Non-proportional patterns of preMN connectivity may allow wing steering MNs to be recruited in different combinations, or with different relative timing. Prior electrophysiological (Balint and Dickinson, 2001; Heide and Götz, 1996; Tu and Dickinson, 1996) and biomechanical (Tu and Dickinson, 1994) studies posit that the action of direct wing MNs are regulated not just by firing frequency, but also by firing phase within the wingbeat cycle, a mechanism that allows a single motor neuron to regulate wing motion with great precision. The connectivity of wing preMNs may provide a basis for flexibly tuning the recruitment order of MNs within a module, whereas the proportional connectivity of leg preMNs provides a basis for stereotyped but fine-scale control of joint force production.

The precision of proportional preMN connectivity onto leg MNs is striking, especially given the fact that leg MN dendrites are spatially intermingled within the leg neuropil (Azevedo et al., 2022). During development, each preMN must find its target module and establish the appropriate number of synapses in proportion to MN size (**Extended Data Figure 5a-c**) (Mendell and Henneman, 1971). To shed light on this question, we next analyze how the structure of motor modules is related to preMN developmental origin.

### PreMNs with diverse developmental lineages contact each motor module

The neurons that make up the *Drosophila* nervous system develop from less than 40 stem cell hemilineages (Shepherd et al., 2019). Neurons within a hemilineage tend to enter the neuropil through common primary neurite tracts, and develop diverse but identifiable morphologies (Lacin et al., 2019; Marin et al., 2023; Truman et al., 2004). We identified the putative hemilineage of local and intersegmental preMNs by comparing stereotyped morphological features to hemilineage-specific genetic driver lines (Harris, 2012; Harris et al., 2015). We also validated our hemilineage identification against a parallel effort in the male VNC connectome (**Figure 5a** and **Extended Data Figure 7**; Marin et al., 2023). We found that some hemilineages contain preMNs that predominantly synapse on either leg or wing MNs (**Figure 5b**). Notably, many different motor modules are targeted by individual local and intersegmental preMNs that that develop from the same hemilineage. Conversely, each motor module also receives synaptic input from preMNs from multiple different hemilineages. In other words, our results demonstrate that a preMN’s preference for a particular module is not strictly determined by its developmental origin.

**Figure 5.**
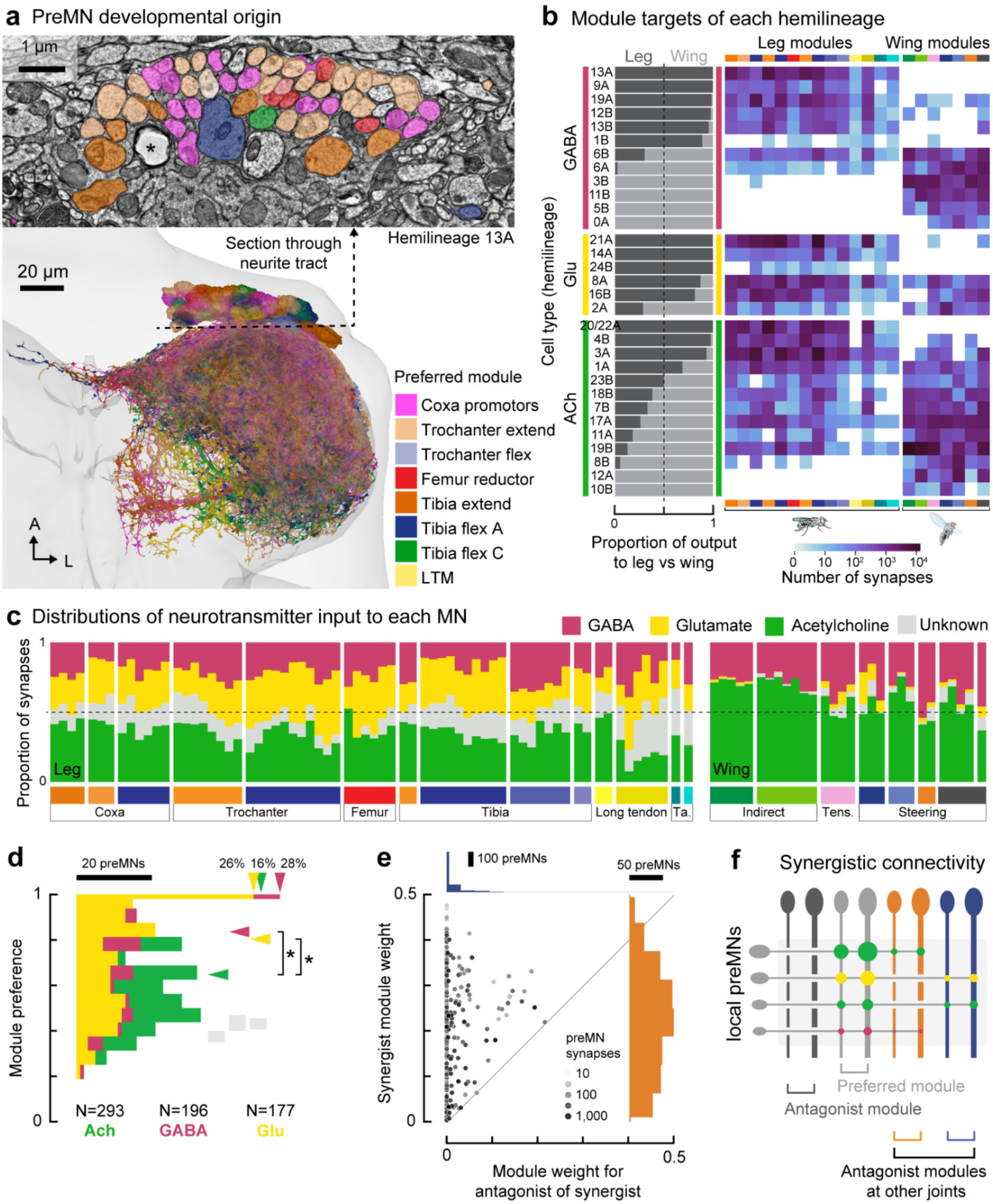
PreMNs with diverse developmental lineages synapse on MNs in each motor module. **a,** PreMNs from an example hemilineage (13A, GABAergic), colored according to their preferred leg motor module. **b,** Left: proportion of the total synapses from all local and intersegmental preMNs in each hemilineage onto leg MNs vs. wing MNs. Right: total synapses onto MNs in each leg or wing module. **c,** Proportion of synaptic input to MNs from cholinergic (green), glutamatergic (yellow), GABAergic (pink), and unidentified (light gray), hemilineages. Only local and intersegmental preMNs are included in this analysis (see Methods). **d,** Module preference of local preMNs, by hemilineage neurotransmitter. Horizontal arrows indicate the median module preference (p=0.0013, Kruskal-Wallis non-parametric one-way ANOVA. Conover’s post-hoc pairwise test with Holm correction for multiple comparisons: * - p=0.002). Gray rectangles indicate the range of medians when preMN connections are shuffled. Vertical arrows show the percentage of neurons that target a single module. **e,** Module weights for synergist module (y-axis, orange histogram), vs. the antagonist of the synergist module (x-axis, blue histogram) (See Methods). Dot hues indicate total synapses onto all MNs. **f,** Schematic of preMNs targeting synergist module while avoiding antagonist modules.

The majority of *Drosophila* neurons release one of three primary neurotransmitters (Allen et al., 2020; Li et al., 2022): acetylcholine, GABA, or glutamate. In the fly central nervous system (CNS), acetylcholine is typically excitatory, acting via nicotinic acetylcholine receptors (nAChR), while GABA is typically inhibitory, acting via GABA_A_ receptors (Allen et al., 2020; Gowda et al., 2018; Lees et al., 2014). Glutamate is excitatory at the fly neuromuscular junction, acting on ionotropic glutamate receptors (GluRs), but can be inhibitory in the CNS, acting on the glutamate-gated chloride channel, GluCl (Liu and Wilson, 2013). GluCl is the most highly expressed glutamate receptor in MNs, although they also express lower levels of GluRIA and GluRIB (Allen et al., 2020).

Most VNC neurons within a hemilineage release the same primary neurotransmitter (Lacin et al., 2019), although exceptions may exist (Marin et al., 2023). We therefore inferred the neurotransmitter of each local and intersegmental preMNs with an identifiable hemilineage (1830 of 2115 = 86% of neurons; 89.9% of synapses). We found that each MN receives a consistent ratio of input from preMNs with different putative neurotransmitters, and that this mix differs across leg and wing systems, with leg MNs receiving more input from putative glutamatergic preMNs (**Figure 5c**). The consistency of this ratio is striking, considering that leg and wing MNs differ up to 40-fold in size and total synaptic input, and that preMNs develop from different lineages and vary up to 100-fold in their total number of output synapses.

Finally, we asked whether local leg preMNs with different putative neurotransmitter identities form different patterns of connectivity with MNs. We found that the module preference value is lower for cholinergic preMNs than for GABAergic or glutamatergic preMNs (**Figure 5d**), indicating that cholinergic preMNs tend to make more synapses onto MNs in non-preferred modules. We then investigated the specificity of these connections outside the preferred module. As described above, the module weight for the antagonist MN is extremely low. For example, the local preMNs that synapse onto trochanter flexor MNs do not synapse on the trochanter extensor MNs and vice versa. Instead, cholinergic leg preMNs tend to synapse onto their preferred module and a second module that controls a different joint, establishing a potential motor synergy. Furthermore, when that secondary module itself has an antagonist, preMNs avoid synapsing onto that antagonist (**Figure 5e**). For instance, if a preMN preferentially synapses on MNs in the Tibia Flex A module as well as trochanter flexor MNs (synergist), then it does not synapse on trochanter extensor MNs (antagonist of the synergist, **Figure 5f**). Glutamatergic and GABAergic preMNs follow this pattern as well (**Figure 5e-f**). Thus, local leg preMNs make stereotyped synaptic connections within their preferred leg modules (**Figure 4b, c, h**), and form precise connectivity patterns across synergistic modules.

## Discussion

Using connectomics, we reconstructed and analyzed the structure of premotor neural circuits that control the fly leg and wing. Our analyses suggest that local VNC interneurons play a central role in integrating descending motor commands with proprioceptive feedback to coordinate MN activity. The connectome is a static anatomical wiring diagram, but the structure of premotor circuits leads to innumerable testable hypotheses for future research, such as specialized functions of motor modules and mechanisms of neural circuit development.

Our analyses also reveal an unexpected level of complexity in the *Drosophila* motor system. We found that a typical fly MN receives thousands of synapses from hundreds of presynaptic preMNs. This number is on par with the scale of synaptic integration in pyramidal cells of the rodent cortex (Schneider-Mizell et al., 2023), though ten times fewer synapses than cat MNs (16,000 - 140,000; (Kernell and Zwaagstra, 1989; Örnung et al., 1998; Ulfhake and Cullheim, 1988), and an order of magnitude more than larval zebrafish MNs (Svara et al., 2018). Vertebrates also exhibit variation in synapse size (Holler et al., 2021), which have not been observed in insects. Thus, synapse counts are thought to be a reasonable estimate of synaptic weights in *Drosophila* (Barnes et al., 2022). Due to technical constraints, we focused on reconstructing and analyzing premotor circuits from a single female fly. However, a parallel effort to reconstruct the VNC connectome of a male fly found similar connectivity patterns, such as descending neurons accounting for ∼10% of premotor input (Cheong et al., 2024). In the future, quantifying how premotor network connectivity varies across individual flies will provide insight into the degree to which these patterns are plastic or hard-wired.

### Modularity of VNC premotor networks

Our analysis of premotor connectivity showed that leg and wing MNs cluster into motor modules formed by patterns of synaptic input from common presynaptic partners. These modules are largely shaped by the synaptic weights of local preMNs. Most modules contain MNs that share clear biomechanical functions. For example, leg modules contain MNs that innervate either the same muscle, or multiple muscles with origins and tendon insertion points that create torque in the same direction (Azevedo et al., 2022; Miller, 1950; Soler et al., 2004). The leg module structure is consistent with the organization of canonical motor pools in other species, such as the gastrocnemius and soleus extensor muscles in vertebrates (Mcphedran et al., 1965; Wuerker et al., 1965). The structures of modules in the wing motor system are different. One pair of wing modules are composed of the MNs that innervate the large, asynchronous indirect power muscles that contract the thorax. Other wing modules are composed of 2-4 synchronous direct wing MNs that control multiple sclerites of the wing hinge for steering. The structure of these steering modules suggests how groups of muscles act via the wing hinge to create subtle 3D deviations in wing trajectory. Some modules align with experimental results and theoretical predictions from past literature (Heide, 1983; Lindsay et al., 2017; Melis et al., 2024; O’Sullivan et al., 2018; Whitehead et al., 2022), while others offer new predictions. For example, we predict that steering module “B”, which contains MNs i1, i2, and b3, is recruited to unilaterally decrease wing stroke amplitude. This prediction is consistent with EMG recordings correlating i1 muscle activity with ipsilateral turns (Heide, 1975), but remains to be tested using causal perturbations.

We found that most preMNs have a strong module preference, in that they make biased synaptic connections onto MNs within the same module. PreMNs within the same developmental lineage also have strong preferences for different modules, consistent with past work in larval *Drosophila* (Mark et al., 2021). A proposed mechanism for wiring precision between preMNs and MNs is that the stereotyped dendritic morphology of MNs that innervate the same muscle allows sequentially born preMNs to find anatomically similar MNs (Baek and Mann, 2009; Brierley et al., 2012; Enriquez et al., 2015; Guan et al., 2022; Mark et al., 2021). In the adult fly, however, the dendrites of MNs in different motor modules are highly intermingled within the neuropil (Azevedo et al., 2022; Balaskas et al., 2019). This lack of spatial topography suggests that spatial location alone is not sufficient for ensuring the wiring precision between preMNs and MNs that we observe in the connectome. Overall, our analyses raise a number of intriguing developmental questions which cannot be answered using connectomics alone, such as what molecular signals cause preMNs from the same hemilineage to target different postsynaptic MNs.

### Specializations of premotor networks controlling limbs with distinct biomechanics

The premotor connectivity matrices for wing and leg MNs provided an opportunity to compare how neural circuits are structured to control musculoskeletal systems with distinct biomechanical constraints. Shared presynaptic input provides a mechanism to control correlations in the firing rates of functionally related MNs. If we define module activity as the vector, **r**(*t*), of firing rates, *r*(*t*), of the MNs in a module (Marshall et al., 2022), then firing rate correlations constrain **r**(*t*), relative to a system in which MNs are independently driven (Henneman et al., 1974). Conceptually, correlated firing rates reduce the entropy of **r**(*t*), i.e., some module activity states cannot occur (Cover and Thomas, 2012). We propose that preMN input, particularly input from local preMNs, may be structured so that the entropy of the module activity matches the biomechanical constraints of the target musculoskeletal elements.

At one extreme, the MNs in the two flight power modules receive nearly the same synaptic weights from shared presynaptic partners. During flight, DLM and DVM MNs all spike at ∼6 Hz (Harcombe and Wyman, 1977; Koenig and Ikeda, 1983). The instantaneous firing rates of DLMs are also desynchronized in a pattern called a “splay state”, such that the DLMs spike one at a time in a regular, repeated pattern (Hürkey et al., 2023). Although their timing is offset, the DLM MNs always spike once within each splay state cycle, which controls the calcium levels in the asynchronously contracting power muscles (Gordon and Dickinson, 2006). Our results suggest that the strong similarity of premotor input to all indirect MNs drives the low frequency central pattern generator that governs the frequency, but not the relative spike timing, of the musculoskeletal flight oscillator.

At the other extreme, clustering the cosine similarity of direct wing MNs revealed modules that arise from preMNs with variable, non-proportional synaptic weights onto the 2-4 MNs in each steering module. In contrast to the indirect muscles, the timing of steering muscle activation is not thought to arise from a central pattern generator (Heide, 1983). We propose that such patterns allow preMNs to vary the correlations between the MN firing rates, allowing **r**(*t*) to range throughout module activity space. We further hypothesize that the advantage of shared synaptic input—relative to independent input to each MN—is the ability to control the precise timing of MN spikes and steering muscle contractions. If each steering muscle has unique effects on the wing’s trajectory (Melis et al., 2024), and these effects depend on the phase of firing within the wingstroke (Heide, 1983; Tu and Dickinson, 1994), then it would be advantageous for preMNs to finely control spike timing while taking advantage of the full module activity space.

Finally, the structure of synaptic input to leg modules reflects correlations in MN firing rates in the form of a recruitment hierarchy, consistent with long-standing predictions of the size principle (Henneman et al., 1974, 1965b). Leg modules arise from the connectivity of excitatory and inhibitory local preMNs that make one-to-all connections onto leg MNs, suggesting that all MNs within a module are collectively excited or inhibited by changes in the activity of each preMN. Each preMN tends to make more synapses onto larger MNs, in proportion to the total synaptic input to that MN. The structure we found is fundamentally different from the connectivity of premotor circuits that control MN recruitment and swimming speed in adult zebrafish, where dedicated pools of V2a premotor neurons synapse onto either slow or fast MNs (Pallucchi et al., 2024; Song et al., 2020). This pattern of roughly equal synaptic weights highlights the need for a reciprocal gradient of excitability to ensure that leg MNs are recruited in the correct order. Patterns of voltage- and ligand-gated ion channel expression likely complement the electrotonic properties of MN morphology to create nonlinear excitable membranes that enforce the rank order of recruitment (Binder et al., 2020, 1983). This result also provides a cautionary tale when interpreting connectomes: without knowledge of MN activity in behaving flies (Azevedo et al., 2020; Sasaki and Burrows, 1998), we would have predicted that MNs that receive the most synaptic input are the most active, when in fact they are rarely recruited.

Past work argues that constraining module activity with a recruitment hierarchy simplifies the problem of how the nervous system controls motor units with a range of neuromuscular properties (Hodson-Tole and Wakeling, 2009). However, violations of the recruitment hierarchy have been observed in many species, particularly during rapid, oscillating movements (Azevedo et al., 2020; Desmedt and Godaux, 1981; Menelaou et al., 2022; Smith et al., 1980). Recent work in primates demonstrated flexible relationships between MN firing rates that changed with the demands of a motor task (Marshall et al., 2022). Integration of common preMN inputs may also vary across MNs of different sizes, thus influencing the trajectory of the module activity. MNs with large dendritic fields, high conductances, and faster time constants may precisely integrate rapidly oscillating excitatory and inhibitory inputs, whereas smaller neurons may be both more sensitive to inhibition and slower to depolarize (Rall, 1959). We found that many intersegmental and descending neurons target specific motor modules, but with less proportional structure than local preMNs. These pathways may evoke flexible correlations in MN firing rates, as they do in primates (Marshall et al., 2022). Thus, we propose that the leg premotor network balances two seemingly opposed demands: reducing the dimensionality of local control within motor modules while maintaining the capacity to flexibly recruit MNs within a module to achieve specific motor tasks.

## Acknowledgements

This work was supported by a Searle Scholar Award, a Klingenstein-Simons Fellowship, a Pew Biomedical Scholar Award, a McKnight Scholar Award, a Sloan Research Fellowship, the New York Stem Cell Foundation, and a UW Innovation Award to JCT; a Genise Goldenson Award to WAL; NIH U19NS104655 to JCT and MD; NIH R01MH117808 to JCT, WAL, and HSS. JCT is a New York Stem Cell Foundation – Robertson Investigator. We thank Jim Truman, David Shepherd, and Elizabeth Marin for assistance with hemilineage identification. We thank Haluk Lacin, Lisa Marin, Greg Jefferis, Gwyneth Card for helpful discussions, and for their lab’s contributions to proofreading in the FANC dataset, particularly Katharina Eichler, Paul Brooks and Gregory Jefferis for sharing comprehensive proofreading and annotation of DNs in the FANC dataset. We thank members of the Tuthill and Dickinson Labs, Sama Ahmed, Bing Brunton, and Jim Truman for comments on the manuscript.

---

**Extended Data Figure 1.**
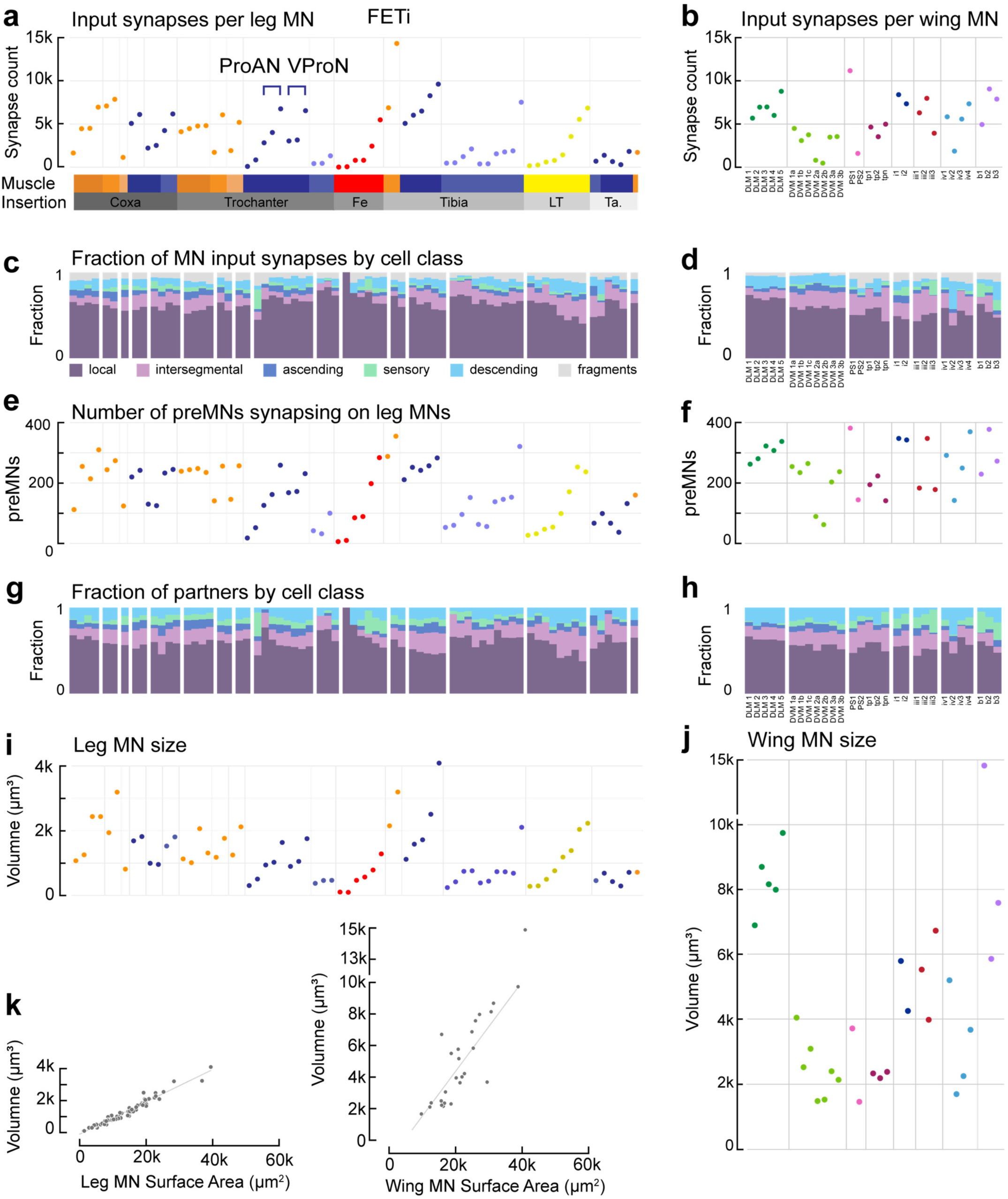
Detailed properties of individual leg and wing MNs. a, Number of input synapses on each leg MN. MNs are ordered by the muscle they innervate, from proximal coxa muscles in the thorax to distal tarsus muscles located in the tibia. **b,** Number of input synapses on each wing MN. Indirect MNs are shown first, direct MNs are ordered according to sclerite. **c,** Fraction of synapses on each leg MN broken down by cell class (see Methods). **d,** Fraction of synapses on each wing MN broken down by cell class. **e,** Number of preMNs presynaptic to each leg MN. **f,** Number of preMNs presynaptic to each wing MN. **g, h,** Fraction of presynaptic partners from each cell class. Presynaptic partners include proofread neurons only, so fragments are not included. **i**, **j**, MN volume. **k,** MN volume vs. surface area for leg MNs (left) and wing MNs (right). Wing MNs tend to have thicker neurites, explaining the steeper relationship. The thick b1 wing steering MN is the outlier.

**Extended Data Figure 2.**
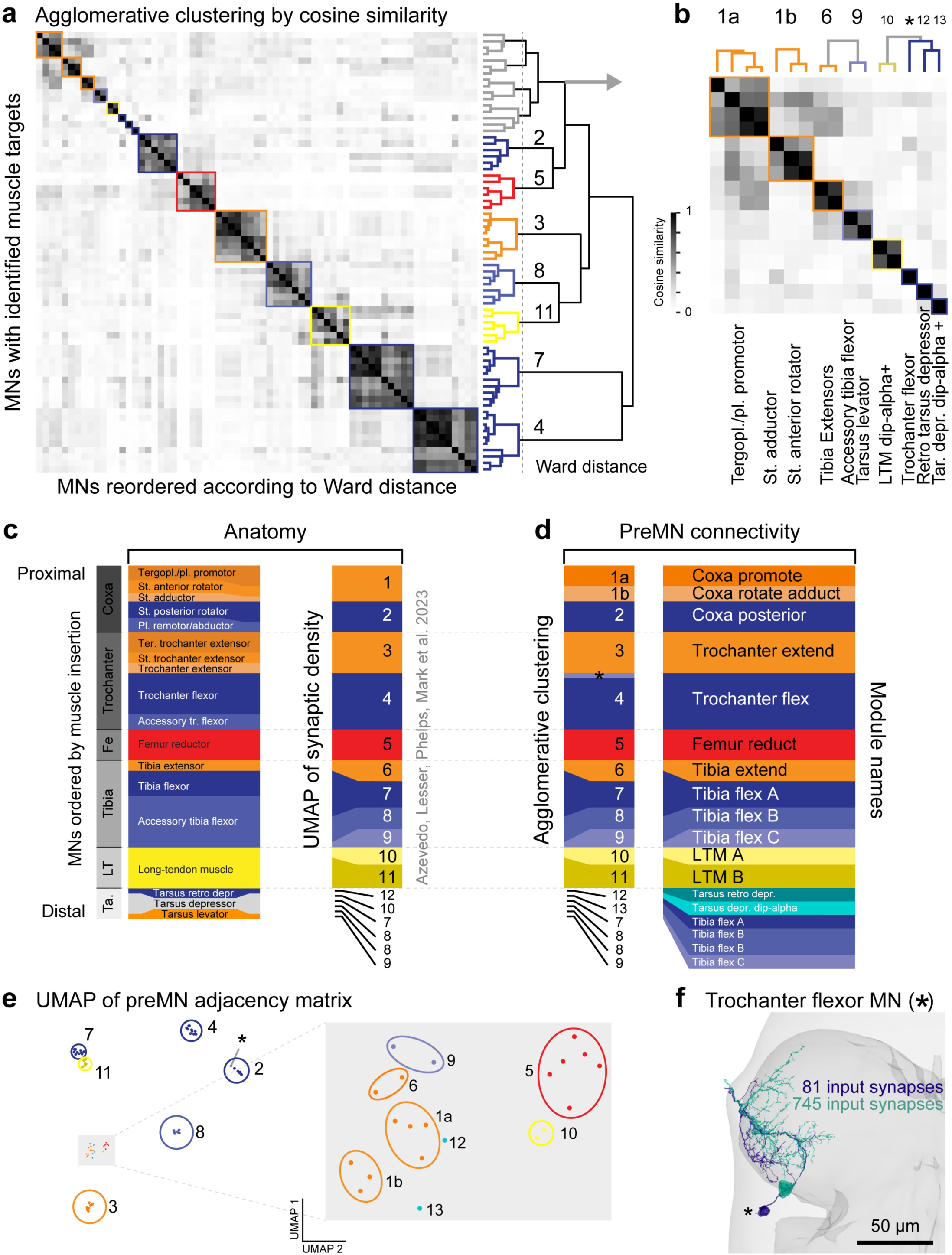
Agglomerative hierarchical clustering of leg MNs according to premotor connectivity. a, Hierarchical clustering of MNs based on the cosine similarity of the columns of the premotor connectivity matrix (Figure 2d). The scipy.cluster AgglomerativeClustering algorithm minimizes the sum of squared distances in each cluster (Ward, scikit-learn). The algorithm identifies seven clear clusters numbered according to the proximal-to-distal origins and insertion points of the innervated muscles (right). **b,** Magnification of seven additional clusters at the top left of the similarity matrix. The muscle targets of MNs in each cluster are indicated below. We remain uncertain about which of four MNs innervate the tergopleural vs. the pleural promotor muscles that insert on the anterior aspect of the coxa (cluster 1a). **c,** In Azevedo et al. 2022, we identified the muscle targets of each MN by comparing anatomical criteria (left). We performed UMAP clustering of the density of input synapses in 3D onto each MN, as an independent, quantitative verification of our anatomical assignments (right). This analysis revealed surprising features that are corroborated by analyzing preMN connectivity here. Specifically, accessory tibia flexor MNs split into 3 distinct clusters, clusters 7, 8, and 9, where the four accessory tibia flexors in cluster 7 clustered with the five main tibia flexor MNs. Additionally, four of the six tarsus MNs clustered with the same groups, one in cluster 7, two in cluster 8, and 1 in cluster 9 (see numbers at bottom). A fifth tarsus MN, the retro depressor MN, clustered on its own (cluster 12). The final tarsus MN clustered with the small LTM MNs in cluster 10; all three are known to express dip-alpha (Venkatasubramanian et al. 2019). **d,** Clusters based on premotor connectivity from **a** and **b**. The differences from the UMAP of synapse density include: promotor MNs of the coxa (cluster 1a) clustered separately from the adductor and rotator MNs (cluster 1b); the tarsus depressor MN (cluster 13) clustered separately from the two dip-alpha-positive LTM MNs (cluster 10, LTM A); a small MN clustered in a separate cluster labeled with an asterisk (*), depicted in **f**. Names for each module are given on the right. **e,** UMAP embedding of the columns of the premotor connectivity matrix (Figure 2d). This clustering does not rely on cosine similarity and largely corroborates the agglomerative clustering. **f,** Two trochanter flexor MNs with somas on the posterior cortex of the neuropil. In total, six MNs have somas on the dorsal cortex. Four of these neurons innervate the sternal posterior rotator muscle. We argued in Azevedo et al. 2022 that the remaining two MNs innervate the trochanter flexor muscle because we observed two axons enter the proximal fibers of the muscle in the X-ray data. The larger of the two MNs (green) clustered with the trochanter flexor MNs according to both the hierarchical clustering and the UMAP embedding. The MN indicated by the asterisk (blue) receives approximately 10X fewer synapses, perhaps explaining why it either clustered by itself (**b)** or with the sternal rotator MNs (**e**). In summary, in the paper, we include the (*) MN with the Trochanter flex MNs.

**Extended Data Figure 3.**
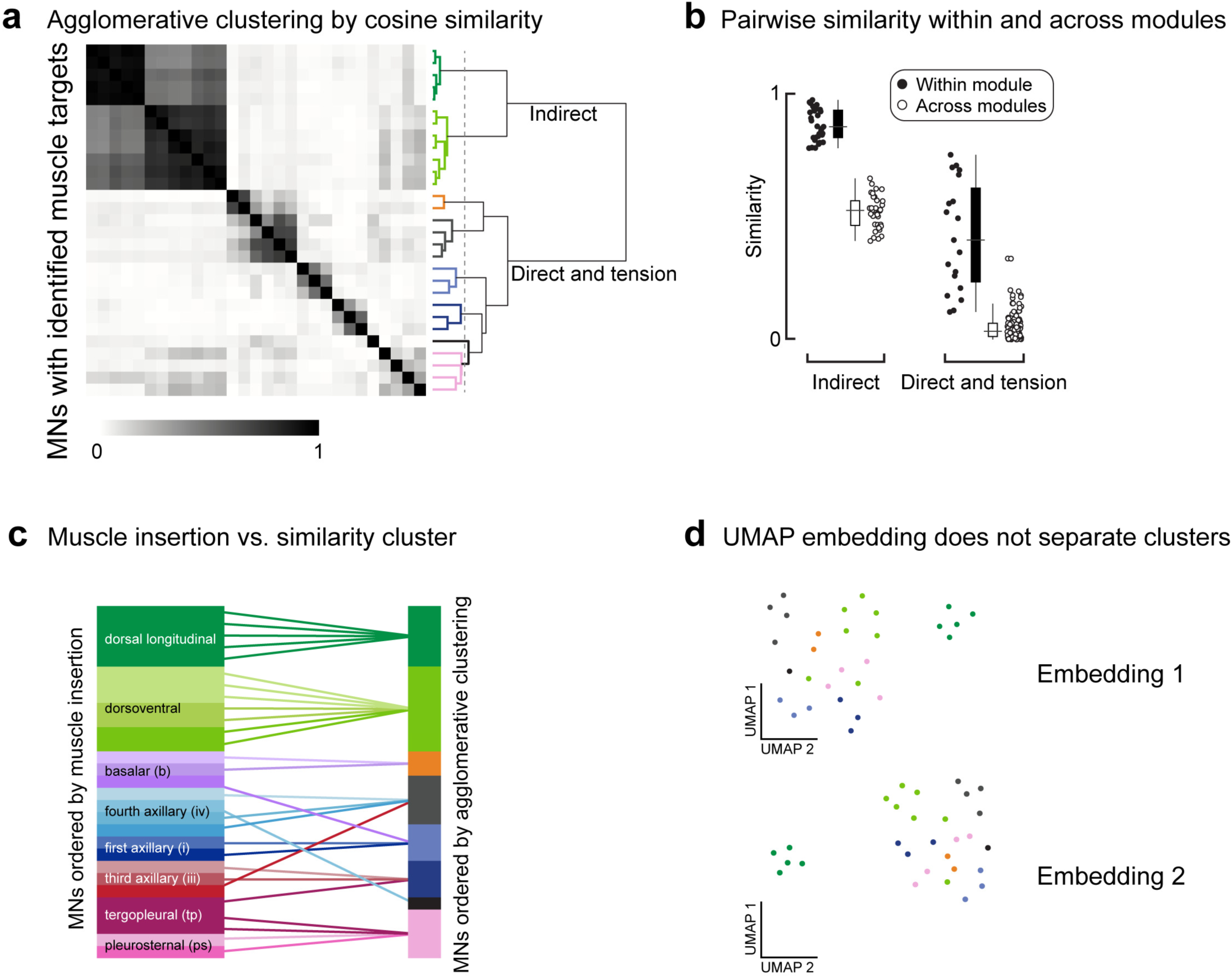
Similarity of wing MNs creates separable modules through agglomerative clustering. a, Cosine similarity matrix for all wing MNs. Axes are symmetric, each row/column is an MN. (Right) The agglomerative clustering dendrogram along with the threshold at which clusters are separable. Colored branches on the dendrogram depict different modules, not muscles. **b,** Similarity scores for each pair of wing MNs. Indirect MNs are separated from direct and tension MNs to better show the distribution of similarity scores of direct and tension MNs. **c,** Schematic showing how ordering by anatomy (left) relates to agglomerative clustering by cosine similarity (right). **d,** UMAP does not separate the wing MNs by connectivity, possibly because their synaptic input weights are not stereotyped (or proportional) from preMNs. Data points are colored post-hoc according to the agglomerative clustering results.

**Extended Data Figure 4.**
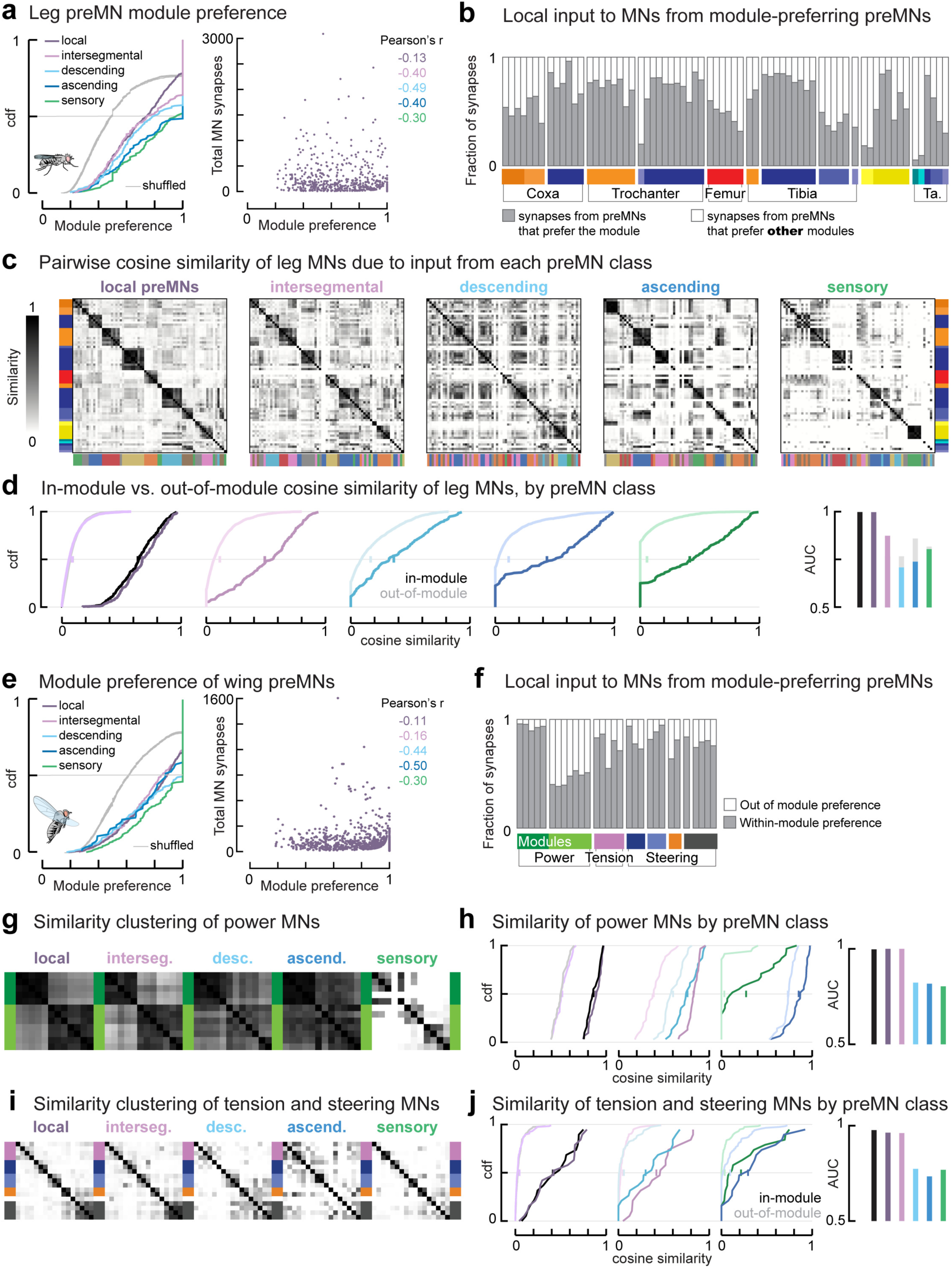
Local PreMNs drive MN cosine similarity and modularity, for wing and leg systems. a, Cumulative density functions (CDFs) of module preference for individual preMNs targeting leg MNs, separated by preMN cell class. Gray indicates ten overlaid example CDFs randomly selected from shuffling the columns of each row of the connectivity matrix. Right, Total MN synapses (y-axis) vs. module preference of local preMNs. Pearson’s r for each cell class is shown, p<10^-13^. PreMNs with fewer MN synapses have a slight tendency to contact a single module. **b**, Fraction of MN input synapses (each bar) from local preMNs that prefer that MN’s module (gray) vs. prefer a different module (white). **c,** Cosine similarity matrices for leg MNs, calculated using synapses from preMNs of each cell class. Modules found in **Extended Data** Figure 2 are shown at left and right. Below each matrix is a color bar indicating clusters found by performing the same agglomerative clustering algorithm on the matrix above. Only local preMN connectivity gives the same clusters as using all preMNs. **d,** CDFs of pairwise similarity of MNs within modules defined in **Extended Data** Figure 2 (dark lines) vs. across modules (light colors). Right, the area under the curve (AUC) measures the overlap of the CDFs, with 0.5 indicating similar CDFs, and 1 indicating complete separation (see Methods). Gray bars show the improvement in the AUC if the clusters shown below the similarity matrices in **c** are used instead. Together, these analyses show that local neurons are responsible for the modularity of MNs. Other classes of preMNs tend to prefer a single module (**a**) but can make select synapses across modules. **e,** Module preference for individual preMNs targeting wing MNs, as in **a**. **f,** Fraction of input synapses on wing MNs from local preMNs that preferentially target each MN’s module (gray) vs a different module (white). **g,** Cosine similarity matrices for indirect (power) MNs, calculated using synapses from preMNs of each cell class. Indirect muscles are divided into two antagonistic modules: dorsal longitudinal muscles (DLMs, dark green) and dorso-ventral muscles (DVMs, light green). They share common input from all cell classes except sensory axons, from which they receive few synapses. **h,** Pairwise similarity of indirect MNs within modules, based on connectivity of each cell class. **i,** Similarity matrices for tension and direct (steering) MNs. **j,** Pairwise similarity of indirect MNs within (dark line) vs. across (light) modules, for each preMN cell class. Colors are indicated in **e**.

**Extended Data Figure 5.**
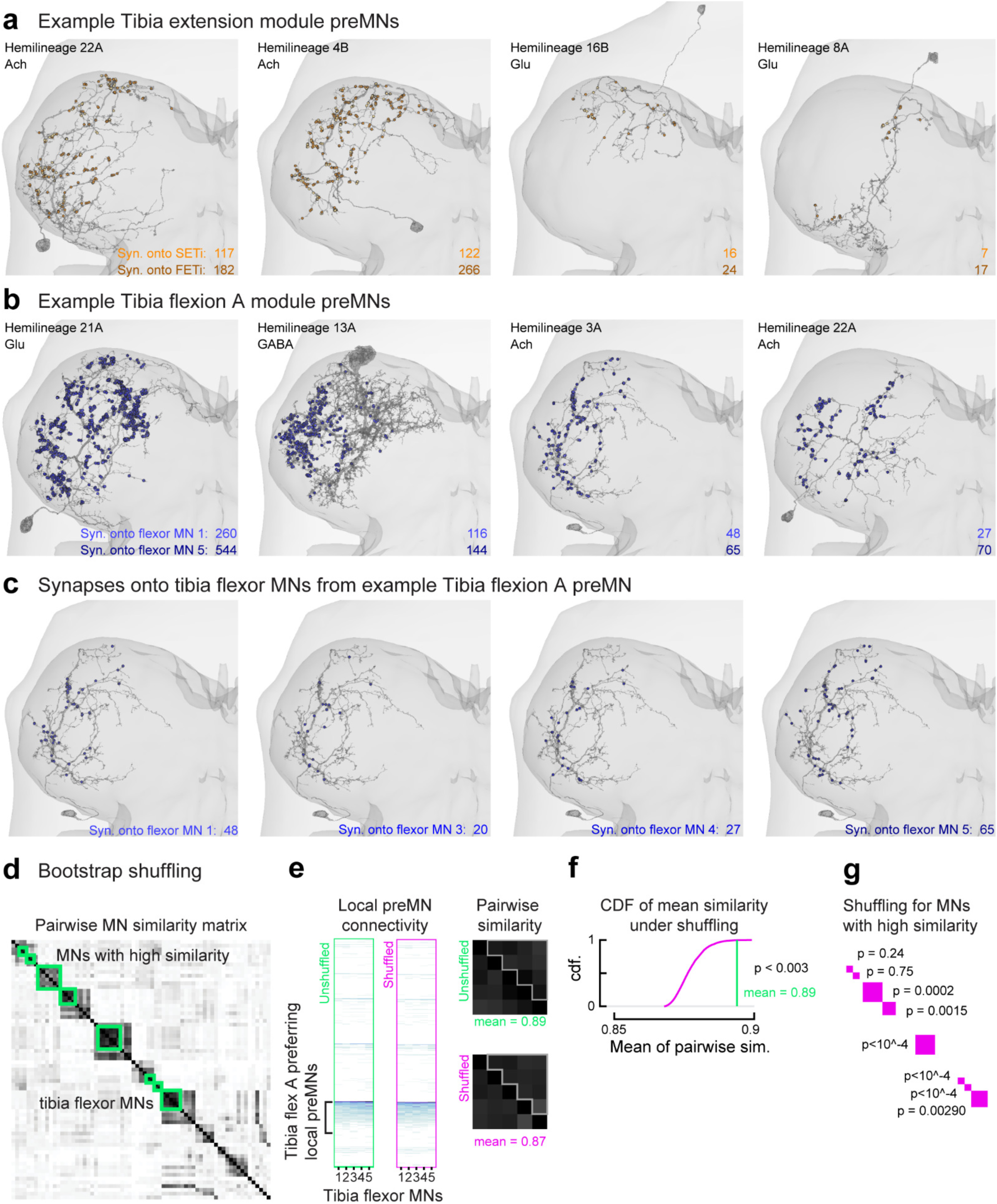
Example leg preMNs, their synapses onto motor modules, and impact of proportional connectivity on MN similarity. a, The location of synapses (spheres) from example preMNs that preferentially synapse onto the SETi (light orange) and FETi (dark orange) tibia extensor MNs. Each preMN has a different morphology and makes more synapses onto FETi than onto SETi. **b,** Example preMNs that preferentially synapse onto the five tibia flexor MNs in the Tibia flex A module (different shades of blue spheres). **c,** A single example preMN from **b**, showing the locations of synapses onto four of the five tibia flexor MNs in the module. The preMN makes more synapses onto the largest neuron, with extra synapses distributed throughout the processes. **d,** Bootstrap shuffling of module connectivity (see Methods for details). This analysis is similar to Figure 4h, but for only the largest neurons with the highest similarity (green squares), where high MN similarity reflects proportional preMN weights onto each MN in the module, as in **a-c**. **e,** Left, the unshuffled synapse counts from all local preMNs onto the Tibia Flex A MNs; middle, the same matrix with example shuffled synapse counts from the module-targeting preMNs; right, the resulting MN similarity matrices, highlighting the pairwise similarities in the upper triangle. **f,** The cumulative probability density function (cdf) of the mean pairwise MN similarity for N=10,000 shuffling repeats, compared to the actual mean. The actual mean is larger than 99.7% of the shuffled instances. **g,** The bootstrap p-value for the regions of high MN similarity. The high p-values indicate pairs of neurons with small differences in their total synaptic input, such that shuffling the proportional synapses does not disrupt a large difference like exists for the FETi and SETi in **a**.

**Extended Data Figure 6.**
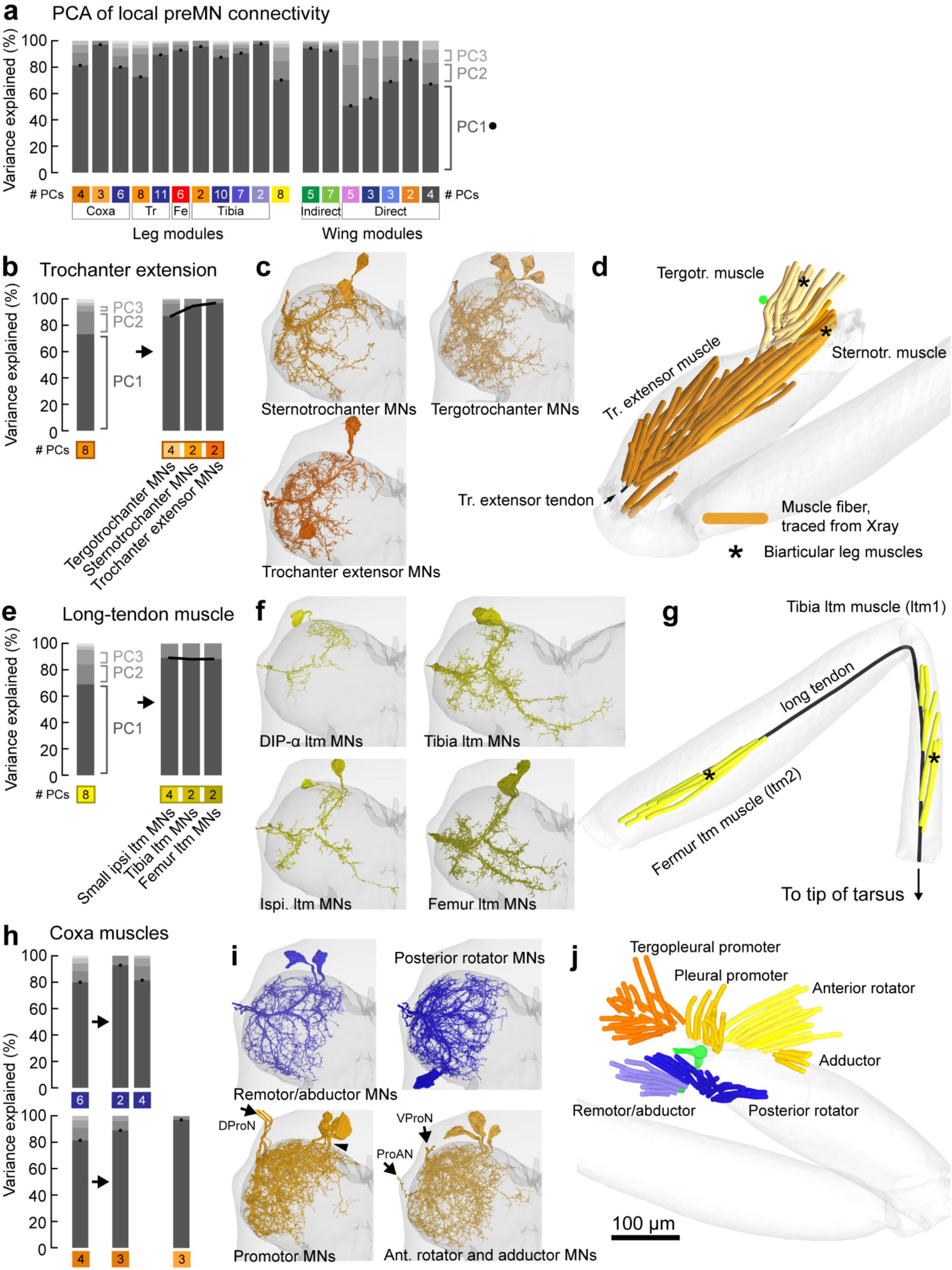
Leg modules that include biarticular muscles have more variable module connectivity. a, Principal Components Analysis of the connections between local preMNs and their preferred motor module. Dots and dark gray bars indicate the percent of the module connectivity variance that is explained by a single principal component, for each leg and wing module. The first principal component was sufficient to explain the majority (>80%) of the variance within most leg motor modules. The percentages of variance explained within modules for the wing power MNs were also high (>90%). For both leg modules and the modules of the wing power muscles, the only significant dimension of variation (captured by the first component alone) was simply the overall preMN output onto the modules. By contrast, the first principal component explained only 63%, 55%, 70%, 87%, and 67% of the variance for the tension, steering A, steering B, C and D modules, respectively. This analysis supports our observation that wing module connectivity is not proportional, from Figure 4h. Here, we further dissect the PCA analysis to show that leg modules that are composed of motor units with more biomechanical diversity have more variable connectivity, in support of our conclusions about the differences between leg and wing connectivity. **b**, The first principal component for the Trochanter extend module explains less of the variance than for most other leg modules. If the module is separated according to muscle target, the first PC explains more of the variance in the connectivity onto MNs targeting each muscle. **c,** MNs innervating the sternotrochanter, tergotrochanter, and trochanter extensor muscles. **d,** Individual traced muscle fibers from the X-ray tomography image volume. The Trochanter extend module contains the biarticular tergotrochanter muscle, which originates at the dorsal thoracic cuticle, crosses the body-coxa joint and extends the trochanter; and the biarticular sternotrochanter muscle, which originates on the ventral thoracic cuticle, crosses the body-coxa joint and extends the trochanter (Azevedo et al. 2022). Note, we adopted the muscle nomenclature from the literature (Miller, 1950). All three muscles insert on the same tendon, so an alternative naming scheme would call these three parts of the same muscle, like the three parts of the human triceps brachii muscle. **e,** When MNs innervating the biarticular long tendon muscle (LTM) are separated by anatomy, the first PC explains more of the connectivity onto MNs targeting each muscle. **f,** Four groups of LTM neurons. The DIP-*ɑ* LTM MNs are small, lack a medial projection, express DIP-*ɑ*, and one targets the femur LTM while the other targets the tibia (Venkatasubramanian *et al*., 2019). The specific muscle targets of two other smaller LTM MNs are unknown. **g,** Traced muscle fibers of the LTM. The LTM is composed of two muscles, one in the femur and one in the tibia, that both insert on the long tendon that crosses multiple articulations to insert on the claw at the tip of the tarsus (Radnikow and Bässler, 1991). **h,** Subdivision of coxa modules, for comparison with biarticular modules. Posterior modules (blue), Anterior modules (orange). Breaking the posterior module into MNs innervating the remotor/abductor muscle or the posterior rotator muscles increases the projection onto the first PC. Excluding a single MN from the coxa promotion module (orange) increases the projection onto the first PC. **i,** MNs innervating the coxa muscles in the thorax. Black arrowhead indicates the primary neurite of the excluded coxa promotor MN. **j,** Coxa muscles in the thorax. We are uncertain about how many MNs innervate the tergopleural vs. the pleural promotor muscles, as the axons are not visible in the X-ray tomography images. The excluded promotor MN receives 4,056 total synapses (compared to 1,484, 4,056, and 6,450 for the other promotor MNs) and has high cosine similarity with the rotator and adductor MNs (**Extended Data** Figure 4C). This analysis suggests that the excluded preMN receives input from slightly different sources than the other preMNs. We speculate that perhaps the excluded promotor MN exits the DProN nerve and innervates the pleural promotor, while the others innervate the tergopleural promotor. In short, the leg modules with more variable input, as measured by PCA, include motor units with more complex biomechanics.

**Extended Data Figure 7.**
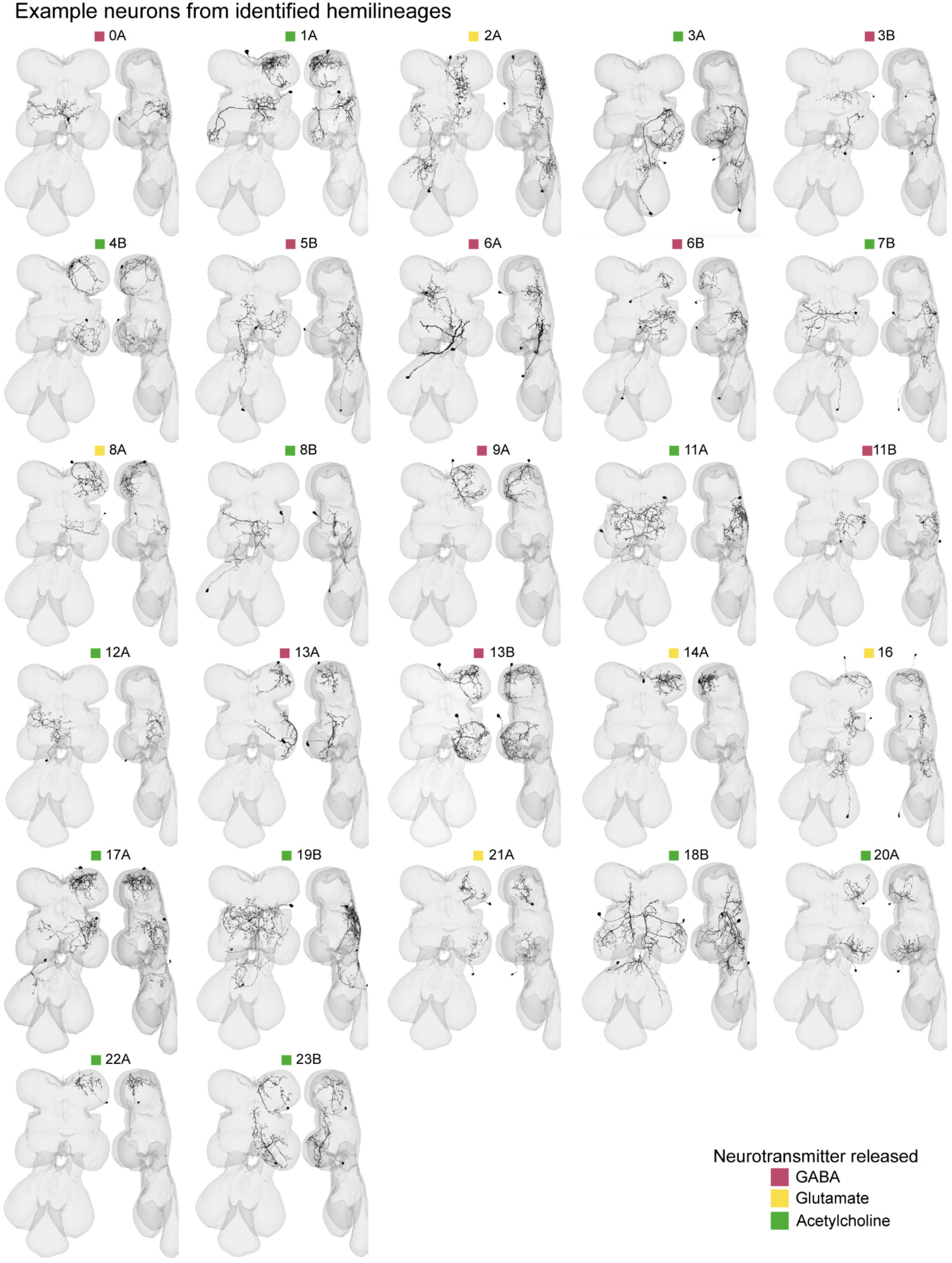
Example neurons from each premotor hemilineage. Example preMNs from each premotor hemilineage. See **Supplemental Table 3** for links to view entire premotor populations of each hemilineage in Neuroglancer, an online tool for viewing connectomics datasets.

**Supplementary Table 1.** Links to motor modules in Neuroglancer. The second columns contains clickable links to view the motor neurons that make up each motor module.

|  |  |
| --- | --- |
| Leg | Coxa promotion<br>Coxa reductor adductor<br>Coxa posterior<br>Trochanter extend<br>Trochanter flex<br>Femur reduct<br>Tibia extend<br>Tibia flex A Tibia flex B<br>Tibia flex C<br>LTM dip-alpha<br>LTM<br>Tarsus retro depressor<br>Tarsus depressor dip-alpha |
| Wing | Dorsal longitudinal<br>Dorsoventral<br>Tension<br>Steering A<br>Steering B<br>Steering C<br>Steering D<br>iv2 |

**Supplementary Table 2.** Links to premotor neurons in Neuroglancer. Every premotor neuron in Figure 2d**,e** organized by their preferred module and cell class.

| **Preferred Module (Leg)** | **Desc.** | **Sensory** | **Asc.** | **Interseg.** | **Local** |
| --- | --- | --- | --- | --- | --- |
| Coxa promotion | link | link | link | link | link |
| Coxa reductor adductor | link | link | link | link | link |
| Coxa posterior | link | link | link | link | link |
| Trochanter extend | link | link | link | link | link |
| Trochanter flex | link | link | link | link | link |
| Femur reduct | link | link | link | link | link |
| Tibia extend | link | link | link | link | link |
| Tibia flex A | link | link | link | link | link |
| Tibia flex B | link | link | link | link | link |
| Tibia flex C | link | link | link | link | link |
| LTM dip-alpha | link | link | link | link | link |
| LTM | link | link | link | link | link |
| Tarsus depressor retro depressor | link | link | link | link | link |
| Tarsus depressor dip-alpha | link | link | link | link | link |
| **Preferred Module (Wing)** |  |  |  |  |  |
| DLM | link | link | link | link | link |
| DVM | link | link | link | link | link |
| Tension | link | link | link | link | link |
| Steering A | link | link | link | link | link |
| Steering B | link | link | link | link | link |
| Steering C | link | link | link | link | link |
| Steering D | link | link | link | link | link |

**Supplementary Table 3.**
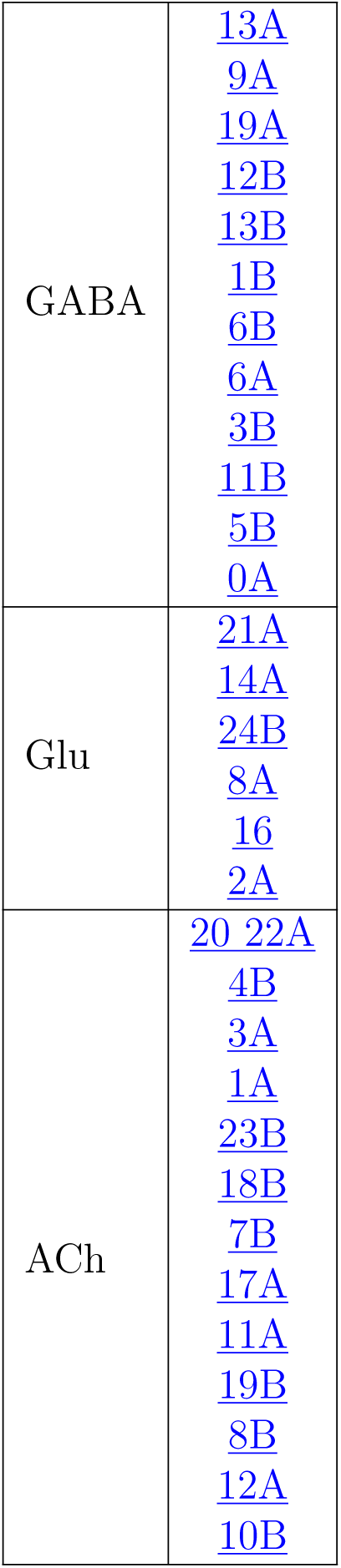
Links to hemilineages in Neuroglancer. Every local and intersegmental premotor neuron that could confidently be assigned to a hemilineage.

## Methods

### Reconstruction of premotor neurons (preMN) anatomy and connectivity

The automated segmentation of the Female Adult Nerve Cord electron microscopy dataset (Phelps et al., 2021), as well as the identification and reconstruction of leg and wing motor neurons (MNs), was described in a companion paper (Azevedo et al., 2022). To manually correct the automated segmentation of premotor neurons (preMNs), we used Google’s collaborative Neuroglancer interface (Maitin-Shepard et al., 2021). We identified all objects in the automated segmentation that were predicted to synapse onto MNs. As a metric of both the quality of the segmentation and the speed of manual proofreading, initially 20% of the pre-synapses were associated with objects that had a soma. We proofread segments until >90% of all input to front left leg and left wing MNs was associated with either a cell body, or an identified descending or sensory process. The remaining inputs were categorized as fragment segments (**Extended Data Figure 1**). We deemed a neuron as “proofread” once its cell body was attached, its full backbone reconstructed, and it had as many branches as could be confidently attached.

Motor neuron volume and surface area were calculated using the statistics of the level 2 cache, which is the graph of “level 2 chunks” in the hierarchy of the ChunkedGraph data structure (Dorkenwald et al., 2022; Schneider-Mizell et al., 2023). Note, the surface area and volume measurements we report do not include the portions of MNs beyond which their axons were cut in the nerve. The computed level 2 cache maintains a representative central point in space, the volume, and the surface area for each level 2 chunk and these statistics are updated when new chunks are created due to proofreading edits. We used the Connectome Annotation Versioning Engine (CAVE) to annotate neurons and keep track of their identities through iterations of proofreading and materializations (Dorkenwald et al., 2023). Somatic segmentations of all motor neurons (downloaded at 68.8 x 68.8 x 45 nm resolution) were cleaned using a heuristic cleaning procedure that removed missing slices of data and incorrectly merged fragments. This procedure has been explained in detail elsewhere (Elabbady et al., 2024). Briefly, a radius of 15 microns surrounding each soma’s center of mass was cut out from the dense segmentation and converted into a binary mask. Subsequent binary dilation in 3D was performed, followed by filling of all holes, and then binary erosion. The resulting binary mask was meshed using marching cubes and connected component analysis was run on the result. For each cell, the somatic surface area and volume measurements were calculated based on the cleaned meshed 15 micron cutout.

### Analysis of MNs controlling the front left leg and the left wing

We described the identification of the muscle targets of the front leg and wing MNs in a companion study(Azevedo et al., 2022), summarized in **Figure 1b**. The leg muscles are named according to the literature (Miller, 1950; Soler et al., 2004).

We focused on the front (prothoracic) leg because 1) we have measured the physiology and neuromechanical properties of MNs innervating the front tibia flexor (Azevedo et al., 2020), 2) the high resolution of our CT volume of the front leg allowed us to count axons entering specific muscles, and 3) T1 contains 69 MNs compared to 62 or 63 for T2 and T3, so we tackled the most challenging task first. We focused on the left T1 because it was better preserved than the right T1(Azevedo et al., 2022).

We analyzed the left wing because the left wing nerve was more intact than the right wing nerve (Azevedo et al., 2022). The MNs in both nerves are intact and fully reconstructed, but the sensory domain of the right wing nerve is damaged.

### Definition of cell classes

We classified all premotor neurons (preMNs) into five groups. Descending neurons had a process in the neck connective and no cell body in the VNC. Ascending neurons had a process in the neck connective and a cell body in the VNC. Sensory neurons had processes entering the VNC from peripheral nerves and no cell body in the VNC. The rest of the neurons were fully contained within the VNC. Leg preMNs were classified as local if all their input synapses were within a bounding box containing the left T1 neuromere, and as intersegmental if they had input synapses outside the bounding box. The bounding box, in pixels, was x = [3000, 38000], y = [90483, 123190], and z = [980, 3858]. Wing preMNs were classified as intersegmental if they had input synapses in any of the six leg neuropils or the abdominal neuropil (e.g., wing preMNs were considered local if they received input from the contralateral wing neuromere, haltere neuromeres, or neck neuromere). We did not split the wing neuropil into right and left sides because some wing MNs cross the midline. All preMNs were manually checked to make sure they were in the correct categories.

In total, we found 271,145 synapses onto leg MNs. We ignored synapses that came from segmented objects that made less than 3 synapses onto MNs, to focus on connections that are less likely to be affected by noise in the synapse detection algorithm(Azevedo et al., 2022), leaving 232,534 synapses (86%). We found 212,190 synapses from the preMN classes above. We also found the following synapses onto leg MNs from sources that fall outside these five cell classes: 4 synapses onto MNs from a neuron with a soma in the VNC and a process in the sensory part of the leg nerve that is not a motor neuron(Phelps et al., 2021); 407 synapses from poorly segmented objects with annotations points in the leg nerve that appear to be merges with glia; and 19,349 synapses from fragments. Finally, we also found 584 output synapses from 37 MNs onto other MNs. Upon inspection, these apparent synapses were all errors; many were at Golgi membranes in the soma, most were dark spots in dendrites, none would have been visually identified as output presynapses with vesicles.

### Definition of motor modules

We previously identified the muscle targets of MNs using anatomical features alone(Azevedo et al., 2022), which we summarize in **Figure 1b**. Here, we used the premotor connectivity matrix to cluster MNs according to shared presynaptic partners (**Figure 3**), agnostic to muscle innervation. For both leg and wing MNs, we calculated the matrix of pairwise cosine similarity of columns of the connectivity matrix (**Extended Data Figure 2, 3**) using the cosine similarity function from the sklearn.metrics module of the sklearn python package(Pedregosa et al., 2011). We then used the AgglomerativeClustering routine from the sklearn python package to merge clusters based on the Ward linkage distance, i.e. minimizing the variance of the merged clusters. The threshold for defining specific clusters differs between the leg and wing data because of differences in network structure and scale.

We chose the cosine similarity measure because it normalizes for total input to each MN, which is the dominant source of variation across MNs. It does not otherwise modify connection weights, thus preserving the quantitative structure of the common input, which we argue is the most salient feature of MN input after normalizing for total input. Cosine similarity is a commonly used technique, including in the connectomics literature, to compare sparse feature vectors, such as the synapse counts across all possible inputs(Li et al., 2020). Alternative similarity measures like the Jaccard index binarize the connection weights(Matsliah et al., 2023), and network measures like weighted stochastic block models(Witvliet et al., 2021) assume probability distributions for connections and weights.

As an alternative to agglomerative clustering, we used the UMAP algorithm to plot the normalized column vectors of the connectivity matrix (L2-norm) on a low dimensional manifold, e.g. from N=1,546-D for the leg matrix to 2-D (**Extended Data Figure 2e**)(McInnes et al., 2020).

To test the robustness of clustering, we compared the distributions of cosine similarity for MNs that clustered together (in-cluster) vs. for MNs in different clusters (out-of-cluster). We quantified the overlap between these distributions using the area-under-the-curve (AUC) metric, as we show in **Extended Data Figure 4d**. The AUC was calculated using the scipy.stats.mannwhitneyu function and dividing by number of comparisons(Virtanen et al., 2020). For leg MNs, the AUC for the assigned clusters is 0.9974, which captures the fact that only a few out-of-cluster comparisons are larger than some in-cluster comparisons. Then, we tested how the AUC changed if each MN was removed from its computed cluster. An increase in the AUC would indicate a better separation between the in-cluster vs. out-of-cluster distributions. The AUC only increased when two small MNs were individually removed from their assigned clusters: the small femur reductor MN that receives a scant 19 total synapses (AUC = 0.9976), and a long-tendon muscle MN that bears enough similarity to the two small long-tendon muscle MNs (dip-alpha positive(Venkatasubramanian et al., 2019)) that removing it from the assigned cluster increased the AUC (0.99785). Thus, except for these two interpretable edge cases, the agglomerative clustering led to the largest difference (largest AUC) between the in-vs. out-cluster distributions of cosine similarity. While we observed a similar overall trend for steering MNs, the AUC was maximized when individual tension MNs were removed from that cluster. This finding corroborates our characterization of tension MNs as bearing little similarity to one another. This gave us confidence that we were not overlooking better ways of clustering, in terms of maximizing cosine similarity within clusters.

Finally, we adopt the synonym “module” to refer to these clusters, based on our finding that most clusters include MNs that innervate muscles with synergistic anatomy, according to our previous identifications.

See **Supplemental Table 1** for links to view MNs grouped by modules in Neuroglancer, an online tool for viewing connectomics datasets.

### Module weight, preferred module, and module preference

We define a module weight as the number of synapses that a preMN makes onto the 1-10 MNs in a module, divided by its total number of synapses onto MNs, expressed as a fraction. Thus, each preMN has a module weight for each module, and likely synapses onto neurons other than MNs. For example, say a preMN makes 100 synapses onto wing MNs, 80 of which are onto different DLM MNs and 20 of which are onto various DVMs. The preMN would have a module weight of 0.8 for the DLM module, 0.2 for the DVM module, and a module weight of 0 for other modules.

We define the module preference as the max over module weight, onto the “preferred module”. Overall, in the leg connectivity matrix, 62.2% of synapses are from preMNs onto their preferred module, and 75.7% for the wing connectivity matrix.

In the Results, we report the module weight for antagonist modules. We considered three groups of antagonist modules: coxa promotor and coxa remotor/adductor modules vs. coxa posterior, trochanter extend vs. trochanter flex, and tibia extend vs. tibia flex A. Together, these modules receive 75% of the synapses from preMNs. We did not include Tibia flex B or C modules because we are uncertain about their role or function. MNs in Tibia flex B and C modules receive more input from preMNs that target other modules, compared to other similarly sized MNs (**Extended Data Figure 4b**).

To analyze the structure of preMN synapses onto non-preferred modules (**Figure 5e**), we found the second highest module weight (after the module preference) for each preMN, which we refer to as a secondary module. If the secondary module had an antagonist, we then compared the module weight for those two modules (see “Module weight, preferred module, and module preference”).

See **Supplemental Table 2** for links to view entire premotor populations that prefer specific modules in Neuroglancer.

### Hypothesis testing by shuffling synapse counts

We compared measurements to null models by computing the same metric when we shuffled the synapse counts in the connectivity matrix, using N=10,000 shuffles. Shuffling the synapse counts preserves both the total number of synapses and the distribution of synapses in a connection (Lynn et al., 2024), but does not preserve information about synapse location. We did not consider randomized connectivity that preserved the total number of synapses but broke apart connections, such as Erdös-Rényi networks or Watts-Strogatz networks(Fornito et al., 2016).

To test whether high cosine similarity (max=0.96, median=0.07) arose stochastically, we computed the maximum and median cosine similarity when we shuffled all synapse counts (max=0.61, median=0.02), just the rows of each column (max=0.54, median=0.03), or just the columns of each row (max=0.82, median=0.09). We conclude that shuffling which preMNs a MN receives input from (columns) lowers the overall distribution of cosine similarity. In contrast, shuffling the synapse counts that each preMN makes onto MNs (rows) never results in the observed high cosine similarity, but it can produce a median cosine similarity comparable to the unshuffled distribution. However, if we fixed the leg modules and shuffled each row, the AUC of the distributions of in-module vs. out-of-module cosine similarity ranged from 0.44 to 0.56, compared to 0.9974 (above, and **Extended Data Figure 4d**).

To test whether high module preference arises stochastically (**Figure 3h**), we shuffled the rows of each column, i.e. the synapse counts that each preMN makes onto MNs. Over N=10,000 shuffles, the median module preference ranged from 0.48 to 0.50, compared to 0.80 for the unshuffled leg MN adjacency matrix. The analogous range is plotted for wing preMNs.

For leg preMNs that contact a module that has an antagonist, the mean module weight for the antagonist module is 0.012, and 84% of preMNs avoid the antagonist altogether. When the preMN synapse counts are shuffled, only 30% of preMNs avoid the antagonist and the mean module weight increases to 0.11.

In **Figure 5d**, we performed the same comparisons but grouped local preMNs that were predicted to use the same neurotransmitters.

In **Figure 4h**, we compared the mean cosine similarity within each module to the shuffled connectivity. Here, we only shuffled the synapse counts that a preMN makes onto the MNs in its preferred module. The probability density function (PDF) of mean cosine similarity for each shuffled matrix was computed by interpolating the cumulative density function (CDF) of mean cosine similarity using linear interpolation from the scipy python package (Virtanen et al., 2020), and differentiating the CDF.

Finally, to test whether proportional connectivity of local preMNs drives high similarity between MNs (**Extended Data Figure 5e**), we considered only the connections from preMNs onto the MNs with the highest similarity, which were the largest MNs in each module. This narrowly tests how often we observe such high MN similarity if we break the tendency of each preMN to make proportionally more synapses onto the largest MNs.

### PCA analysis of module connectivity

We define module connectivity as the matrix of synapse counts onto all of the MNs in a module, from the preMNs that preferentially target that module. We performed PCA on the module connectivity for each module using the scikit-learn PCA routines in python (Pedregosa et al., 2011) (**Extended Data Figure 6**). The number of PCs for each module is the same as the number of MNs in the module. The first PC is the output weights onto MNs in the module that captures the most variance. By definition, the remaining PCs must be orthonormal and together capture the remaining variability in the module connectivity. For this analysis, we combined all of the long-tendon muscle MNs (8 MNs).

### Identification of premotor hemilineages

The assignment of premotor neurons into hemilineages was based on a suite of evidence, the foundation of which are hemilineage-specific split-Gal4 lines created by Jim Truman, Haluk Lacin, and David Shepherd, which they have experimentally validated, both in prior work (Harris et al., 2015; Lacin et al., 2019; Lacin and Truman, 2016) and in manuscripts that are still in preparation. These split-Gal4 lines identified anatomical hallmarks, namely the neurite bundle through which each cell enters the neuropil, that have allowed us and others to determine hemilineage identity. We also consulted with our colleagues working in the male VNC connectome (MANC), particularly Lisa Marin and Greg Jefferis, to ensure continuity of hemilineage identifications across FANC and MANC and to resolve any edge cases. Since the submission of our paper, the MANC group has released a preprint describing the identification of VNC hemilineages (Marin et al., 2023). That effort included the deployment of a convolutional neural network (CNN) that predicts the neurotransmitter at a synapse from the EM images (Eckstein et al., 2020). The CNN predicted that over 80% of neurons in each hemilineage express the same transmitter, and the same transmitter reported by Lacin et al. 2019. One exception, hemilineage 09B, was predicted to produce both cholinergic and glutamatergic neurons roughly equally, but we identified no 09B neurons as preMNs. Of the 2,115 local and intersegmental preMNs for both the leg and wing, 1,830 were matched to a hemilineage (**Extended Data Figure 7**), which accounted for 89.9% of the VNC input to left front leg and left wing MNs.

See **Supplemental Table 3** for links to view entire premotor populations of each hemilineage in Neuroglancer.

## Software and data availability

Data presented in the paper was analyzed from the CAVE materialization version 840 (v840). Annotated connectivity matrices (**Figure 2**) are available as python Pandas data frames (https://pandas.pydata.org/) at the git-hub repository for this paper, https://github.com/tuthill-lab/Lesser_Azevedo_2023. Also available at the repository are scripts to recreate the analyses and figures in the paper, as well as scripts to recreate the connectivity matrices, for users authorized to interact with the CAVEclient. Links to public preMN and MN segmentations are available throughout the text, as well as in **Supplemental Tables 1-3**. All analysis was performed in Python 3.9 using custom code, making extensive use of CAVEclient (https://github.com/seung-lab/CAVEclient) (Dorkenwald et al., 2023) and CloudVolume to interact with data infrastructure, and libraries Matplotlib, Numpy, Pandas, Scikit-learn, Scipy, stats-models and VTK for general computation, machine learning and data visualization. Additional code is available at https://github.com/htem/FANC_auto_recon, providing additional tutorials, documentation for interacting with FANC, and instructions for joining the FANC community.

